# The ultrastructural properties of the endoplasmic reticulum govern microdomain signaling in perisynaptic astrocytic processes

**DOI:** 10.1101/2022.02.28.482292

**Authors:** Audrey Denizot, María Fernanda Veloz Castillo, Pavel Puchenkov, Corrado Calì, Erik De Schutter

## Abstract

Astrocytes are now widely accepted as key regulators of brain function and behavior. Calcium (Ca^2+^) signals in perisynaptic astrocytic processes (PAPs) enable astrocytes to fine-tune neurotransmission at tripartite synapses. As most PAPs are below the diffraction limit, their content in Ca^2+^ stores and the contribution of the latter to astrocytic Ca^2+^ activity is unclear. Here, we reconstruct hippocampal tripartite synapses in 3D from a high resolution electron microscopy (EM) dataset and find that 75% of PAPs contain some endoplasmic reticulum (ER), a major calcium store in astrocytes. The ER in PAPs displays strikingly diverse shapes and intracellular spatial distributions. To investigate the causal relationship between each of these geometrical properties and the spatio-temporal characteristics of Ca^2+^ signals, we implemented an algorithm that generates 3D PAP meshes by altering the distribution of the ER independently from ER and cell shape. Reaction-diffusion simulations in these meshes reveal that astrocyte activity is governed by a complex interplay between the location of Ca^2+^ channels, ER surface-volume ratio and spatial distribution. In particular, our results suggest that ER-PM contact sites can act as local signal amplifiers if equipped with IP_3_R clusters but attenuate PAP Ca^2+^ activity in the absence of clustering. This study sheds new light on the ultrastructural basis of the diverse astrocytic Ca^2+^ microdomain signals and on the mechanisms that regulate neuron-astrocyte signal transmission at tripartite synapses.

**Main points:**

- 6 nm isotropic volume EM reveals that 75% of perisynaptic astrocytic processes (PAPs) contain some ER
- PAPs & ER of the same cell display diverse geometric properties
- Simulations in EM-derived meshes hint that ER geometry governs Ca^2+^ signals in PAPs

**TOC:** 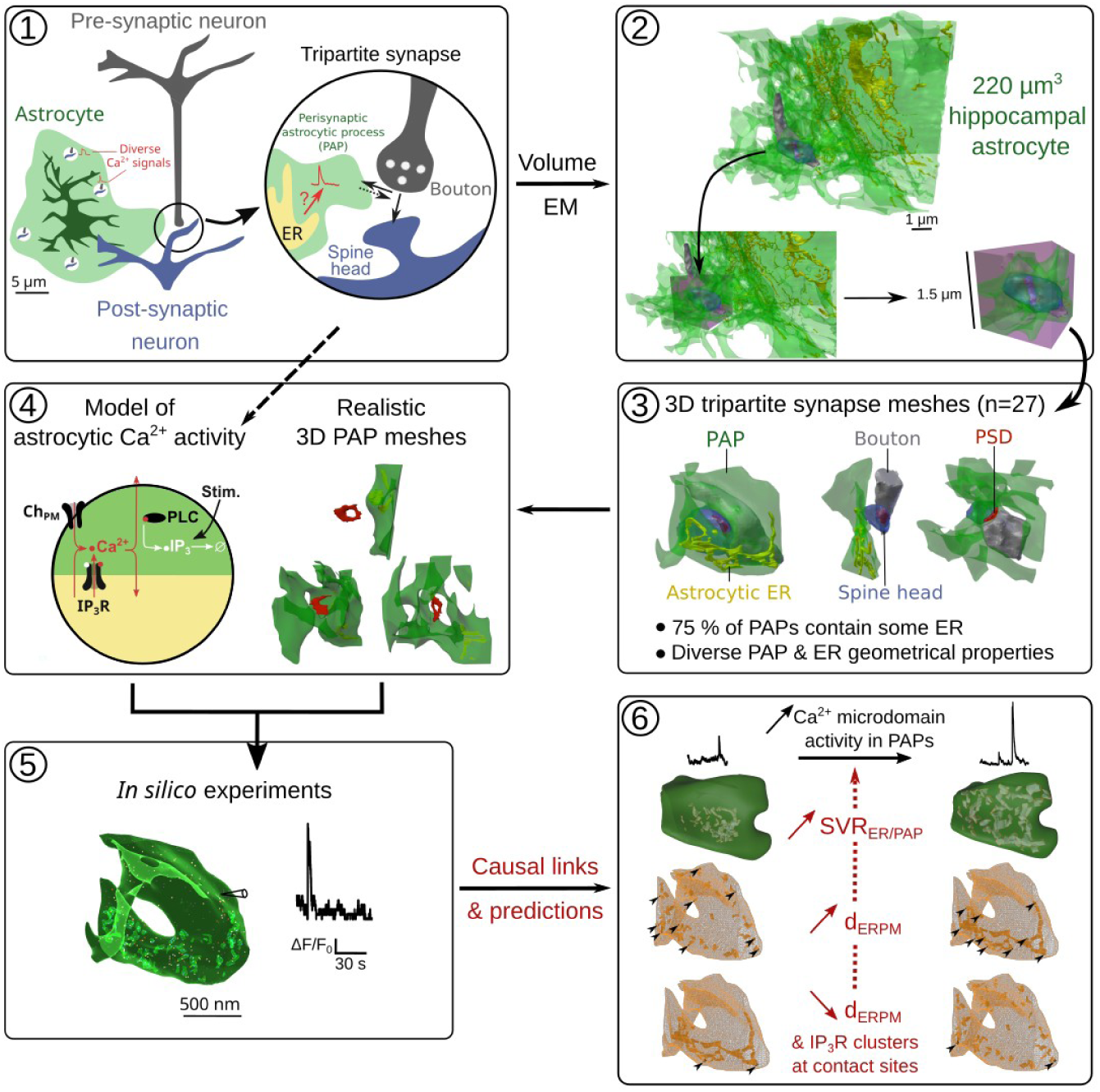

## Introduction

Astrocytes, the most abundant glial cells of the central nervous system, are essential for numerous brain functions and shape behavior [1, 2, 3]. In particular, astrocytes are key modulators of neurotransmission at tripartite synapses [4, 5]. A single astrocyte in the CA1 region of the mouse hippocampus is in contact with hundreds of thousands of synapses simultaneously, at perisynaptic astrocytic processes (PAPs) [6]. Around 75 % of cortical and 65 % of hippocampal synapses are contacted by an astrocytic process [7, 8]. This close contact between astrocytes and neurons allows astrocytes to control various synaptic functions, from glutamate uptake [9], and spillover [10, 11], to synapse homeostasis [12], stability [13], synaptogenesis [14], and neurotransmission [15, 5]. Those synaptic functions are associated with specific local molecular expression in PAPs [16, 17], which changes upon fear conditioning [16]. Importantly, the alteration of the proximity of PAPs to hippocampal synapses of the CA1 region *in vivo* affects neuronal activity and cognitive performance [11]. Conversely, neuronal activity has been shown to induce the remodeling of synaptic coverage by PAPs in various brain regions, both *in vivo* and in acute slices [10, 18, 19, 13, 8, 20, 21, 22]. Together, these results illustrate that PAPs are preferential sites of neuron-astrocyte communication. Although the recent emergence of super-resolution techniques has provided key insights into the properties and functions of PAPs [23, 24], our understanding of PAP physiology and function in live tissue is hindered by their nanoscopic size [25, 26].

Ca^2+^ signals are commonly interpreted as a measure of astrocyte activity, notably in response to neurotransmitter release at synapses [27, 25, 28]. The recent advances in Ca^2+^ imaging approaches have improved the spatio-temporal resolution of Ca^2+^ signals monitored in astrocytes [29, 28]. Strikingly, it revealed that astrocytes in acute slices and *in vivo* exhibit spatially-restricted Ca^2+^ signals, also referred to as hotspots or microdomains, that are stable over time, and which activity varies under physiological conditions such as locomotion or sensory stimulation [30, 31, 32, 33, 34, 35, 36, 37, 38, 39, 40, 41, 42]. Growing evidence supports that PAPs are preferential sites displaying spatially-restricted Ca^2+^ microdomains in response to neurotransmission [40, 41, 43, 44, 30]. As a single astrocyte can contact hundreds of thousands of synapses simultaneously [6], such spatially-restricted Ca^2+^ microdomains might enable the astrocyte to finely tune synaptic transmission at the single synapse level.

mGluR activation on the astrocytic membrane following neurotransmission at glutamatergic synapses results in Ca^2+^ transients mediated by *G_q_*proteins and Ca^2+^ stores such as the endoplasmic reticulum (ER) [29], which can trigger the release of molecules that modulate neurotransmission, gliotransmitters [45, 46, 15, 5]. Most astrocytic Ca^2+^ signals are mediated by the Inositol 3-Phosphate (IP_3_) receptors on the membrane of the endoplasmic reticulum (ER) [47]. Because of their nanoscopic size, the Ca^2+^ pathways involved in microdomain Ca^2+^ signals in PAPs are still unclear. In particular, the presence of ER in PAPs and its involvement in microdomain Ca^2+^ signals at synapses are highly debated. During the last decade, PAPs have been regarded as devoid of ER, with a minimum distance between the synapse and the closest astrocytic ER *>* 0.5 *µ*m [48, 25, 49]. In contrast, recent studies suggest that Ca^2+^ activity in PAPs partly results from Ca^2+^ fluxes from the ER. Notably, inhibiting ER-mediated Ca^2+^ signaling in fine processes results in a decreased number of Ca^2+^ domains (Figure 4N-Q of [32]) and a decreased Ca^2+^ peak frequency [32, 44, 39]. Furthermore, some astrocytic ER has been detected near synapses in recent EM studies [26, 50]. Yet, the geometrical properties of the ER in PAPs and its distribution remain poorly characterized, but could have a strong impact on neuron-astrocyte communication at tripartite synapses.

**Table 1:**
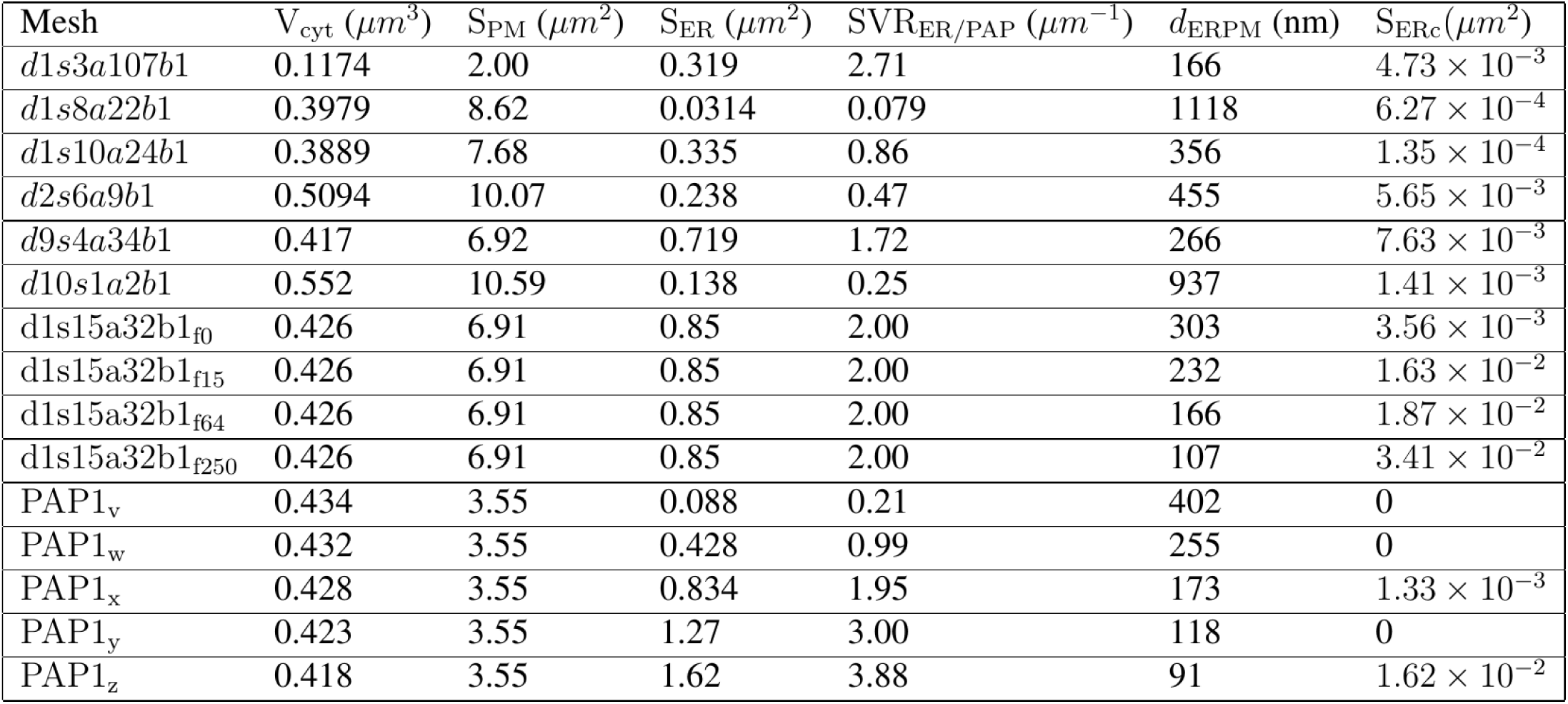
Characteristics of the 3D PAP meshes used in the reaction-diffusion simulations. V_cyt_ is the cytosolic volume, S_PM_ is the plasma membrane surface area, S_ER_ is the ER surface area, SVR_ER*/*PAP_ is the ratio between the ER surface area and the cytosolic volume. d_ERPM_ is the median ER-plasma membrane distance. S_ERc_ is the ER surface area at ER-PM contact sites, i.e. where *d*_ERPM_ ≤ 20 nm . d1s15a32b1_f0_, d1s15a32b1_f21_, d1s15a32b1_f64_ and d1s15a32b1_f250_ refer to meshes from frames 0, 21, 64 and 250 of the d1s15a32b1 PAP mesh presented in Fig. 5.

| Mesh | $V_{\text{cyt}} (\mu m^3)$ | $S_{\text{PM}} (\mu m^2)$ | $S_{\text{ER}} (\mu m^2)$ | $SVR_{\text{ER/PAP}} (\mu m^{-1})$ | $d_{\text{ERPM}} (\text{nm})$ | $S_{\text{ERc}} (\mu m^2)$ |
| --- | --- | --- | --- | --- | --- | --- |
| *d1s3a107b1* | 0.1174 | 2.00 | 0.319 | 2.71 | 166 | $4.73 \times 10^{-3}$ |
| *d1s8a22b1* | 0.3979 | 8.62 | 0.0314 | 0.079 | 1118 | $6.27 \times 10^{-4}$ |
| *d1s10a24b1* | 0.3889 | 7.68 | 0.335 | 0.86 | 356 | $1.35 \times 10^{-4}$ |
| *d2s6a9b1* | 0.5094 | 10.07 | 0.238 | 0.47 | 455 | $5.65 \times 10^{-3}$ |
| *d9s4a34b1* | 0.417 | 6.92 | 0.719 | 1.72 | 266 | $7.63 \times 10^{-3}$ |
| *d10s1a2b1* | 0.552 | 10.59 | 0.138 | 0.25 | 937 | $1.41 \times 10^{-3}$ |
| *d1s15a32b1<sub>f0</sub>* | 0.426 | 6.91 | 0.85 | 2.00 | 303 | $3.56 \times 10^{-3}$ |
| *d1s15a32b1<sub>f15</sub>* | 0.426 | 6.91 | 0.85 | 2.00 | 232 | $1.63 \times 10^{-2}$ |
| *d1s15a32b1<sub>f64</sub>* | 0.426 | 6.91 | 0.85 | 2.00 | 166 | $1.87 \times 10^{-2}$ |
| *d1s15a32b1<sub>f250</sub>* | 0.426 | 6.91 | 0.85 | 2.00 | 107 | $3.41 \times 10^{-2}$ |
| PAP1<sub>v</sub> | 0.434 | 3.55 | 0.088 | 0.21 | 402 | 0 |
| PAP1<sub>w</sub> | 0.432 | 3.55 | 0.428 | 0.99 | 255 | 0 |
| PAP1<sub>x</sub> | 0.428 | 3.55 | 0.834 | 1.95 | 173 | $1.33 \times 10^{-3}$ |
| PAP1<sub>y</sub> | 0.423 | 3.55 | 1.27 | 3.00 | 118 | 0 |
| PAP1<sub>z</sub> | 0.418 | 3.55 | 1.62 | 3.88 | 91 | $1.62 \times 10^{-2}$ |

Recent advances in electron microscopy (EM) enable the resolution of the ultrastructure of astrocytes at an unprecedented spatial resolution. Here, we reconstruct 46 three dimensional meshes of tripartite synapses from a 220 *µm*^3^ hippocampal astrocytic volume from the CA1 stratum radiatum region (6 nm voxel resolution) [51], reconstructed from electron microscopy (EM). Strikingly, we find that 75 % of PAPs in this dataset contain some ER, which can be as close as 72 nm to the post-synaptic density (PSD). Analysis of the geometrical features of these meshes reveal the vast diversity of ER shapes and distributions within PAPs from a single cell. We then used a detailed stochastic reaction-diffusion model of Ca^2+^ signals in PAPs to investigate the mechanistic link between the spatial characteristics of the ER measured in the 3D meshes and the spatio-temporal properties of Ca^2+^ microdomain activity in PAPs. To be able to decipher the effect of ER distribution within the PAP independently from the effect of its shape, we developed an algorithm that automatically creates realistic 3D tetrahedral PAP meshes with various ER distributions based on realistic meshes reconstructed from EM. *In silico* experiments in these meshes reveal that the spatio-temporal properties of Ca^2+^ signals in PAPs are tightly regulated by the intracellular geometry. Together, this study provides new insights into the geometrical properties of hippocampal tripartite synapses and predicts causal links between these features and Ca^2+^ microdomain activity at tripartite synapses.

## 2 Methods

### 2.1 3D reconstruction from electron microscopy

#### 2.1.1 Sample preparation and imaging

The original dataset used in this work (EM stack and 3D reconstructions) was previously published in [51]. The block was a gift from Graham Knott (BioEM imaging facility at EPFL, Lausanne, Switzerland). All procedures were performed according to the Swiss Federal Laws.

One P90 Sprague-Dawley rat was deeply anesthetized with isoflurane and transcardially perfused using 2 % paraformaldehyde and 2.5 % glutaraldehyde in PBS 0.1M. Coronal sections (100 *µ*m) were obtained and washed in cacodylate buffer, followed by a post-fixation using osmium tetroxide and uranyl acetate. Finally, the sections were embedded in Durcupan. Regions of the hippocampus were dissected under a stereoscopic microscope, mounted onto a blank resin slab, and trimmed using an ultramicrotome (Laica Ultracut UC-7). Imaging was performed using an NVision 40 FIB-SEM (Carl Zeiss) with an acceleration voltage of 1.5 kV, a current of 350 pA, and a dwell time of 10 *µ*s/pixel. Serial images were obtained using backscattered electrons and collected at a 6 nm/pixel magnification and a 5 nm milling depth between images.

#### 2.1.2 3D reconstruction and rendering

The serial micrographs were first registered using Multistackreg, a freely available plug-in for Fiji [51]. Then, using those micrographs, we proceeded to the image segmentation and 3D model reconstructions by using TrackEM2 (a plug-in for Fiji) for manual segmentation, and iLastik, for a semi-automated segmentation. The extracted models were then imported into Blender software for visualization and rendering purposes [52].

#### 2.1.3 Extraction of tripartite synapse meshes

For each synapse in contact with the 220 *µm*^3^ astrocytic volume, a cube of edge length 1.5 *µm* (3.375 *µm*^3^) was created and centered at the center of mass of the PSD. All of the elements of the mesh (astrocyte, astrocytic ER, spine and bouton) that were within the cubic volume were isolated using a boolean intersection operator available in Blender, forming what we refer to as a tripartite synapse mesh.

The size of the cube was chosen to be large enough to contain the whole spine and bouton elements while containing a single synapse, taking into consideration that the neuropil is believed to contain around one synapse per micrometer cube. This workflow resulted in the creation of 44 excitatory and 2 inhibitory synapse meshes.

### 2.2 3D mesh manipulation

All 3D mesh manipulations were performed with open-access, open-source software. All 3D meshes used in this study are freely available under the Creative Commons Attribution 4.0 International license at https://zenodo.org/records/17106549.

#### 2.2.1 3D PAP mesh processing for reaction-diffusion simulations

PAP meshes from tripartite synapse meshes were pre-processed using Blender software to be suitable for reaction-diffusion simulations. The workflow is illustrated in Fig. S8. Intersection between ER and PAP membranes was prevented by using a boolean intersection operator. ER was relocated a few nanometers away from the plasma membrane. PAP compartments that were disconnected from the largest PAP volume were deleted. Boolean difference operation was performed between the PAP and ER elements. Non-manifold vertices were repaired. The resulting PAP surface mesh was exported in stl format, which was then converted into a 3D tetrahedral mesh (msh format) using TetWild software [53]. Lastly, the mesh was imported into Gmsh software to be converted into 2.2 ASCII format, supported by the STEPS mesh importer.

#### 2.2.2 Generation of realistic PAP meshes with various ER distributions and constant shape and size

We have implemented a script that generates realistic 3D tetrahedral PAP meshes characterized by various ER locations, constant ER shape and size. The algorithm is written in Python, can be imported in Blender, and is available at https://github.com/adenizot/PAP-ER. The workflow is presented in Fig. 5A. First, all elements of the mesh, i.e. the PAP and the ER, are relocated so that their center of mass are centered at the origin. Then, the ER is split into smaller ER objects using a custom-made function. Briefly, n cubes of a given size are placed along the ER object. Intersection boolean operation is then performed between the ER and each cube, resulting in the creation of n ER objects. ER objects smaller than 30 nm^3^ are deleted. The remaining ER objects are rescaled so that the sum of their surface areas matches the area of the original ER element, measured with the “3D Print” Blender add-on. The number and size of cubes can be altered depending on the size of the original ER and on the mesh characteristics desired. Using Blender’s physics engine, a simulation with m frames is generated, in which ER objects are subject to physical forces that alter their location between each frame. The input of the “RunPhysics” function includes parameters that affect how close objects can get, which can be altered to prevent membrane intersection. Note that parameter values used in the ER splitting and scattering functions should be adjusted depending on the mesh used (see the repository at https://github.com/adenizot/PAP-ER for more details). Examples of frames generated by this workflow applied to d1s15a32b1 PAP mesh are presented in Supplementary movie 3. For each selected frame, the mesh pre-processing steps presented in Fig. S8 are performed automatically, resulting in the export of a surface mesh (stl format). 3D meshing and format conversion can then be performed using TetWild and Gmsh software, as described above. The resulting meshes can be used to perform reaction-diffusion simulations.

#### 2.2.3 Analysis of the geometrical properties of 3D meshes

The volume and surface area of each synaptic element, i.e. the PAP, astrocytic ER, spine head, and bouton, were measured using the Blender add-on “3D Print”. We implemented a Python script that can be imported in Blender 4.3.2. software (https://www.blender.org/) that measures distances between the mesh elements of interest. The code is available at https://github.com/adenizot/PAP-ER. The distance between each vertex of the plasma membrane (PM) of the PAP and the center of mass of the neighboring PSD, and the closest vertices on the bouton and spine head membranes were computed in Blender and stored in a list. Similarly, ER-PSD distance was analyzed by measuring the distance between each vertex of the ER membrane and the center of mass of the PSD. To characterize the distribution of the ER, for each vertex on the PM, the closest ER vertex was detected and its distance to the PM vertex was stored in a list. To compare the distribution of the ER in different 3D meshes, we computed the median of ER-PM distances in each mesh: d_ERPM_. PAP-PSD, PAP-Bouton, PAP-Spine, ER-PSD, and ER-PM distance lists were exported to a text file for analysis and visualization.

### 2.3 Computational modeling

#### 2.3.1 Modeled reactions and computational approach

Astrocytic Ca^2+^ signals in PAPs were simulated using the reaction-diffusion voxel-based model of ER-dependent Ca^2+^ signaling from Denizot and colleagues ([54] Table 2, Fig. 6-7). The kinetic scheme is presented in Figure 3A and Table S2. Parameter values and initial conditions are detailed in Table S3. Briefly, the model describes Ca^2+^ fluxes in and out of the astrocytic cytosol. The opening of IP_3_R channels on the ER membrane triggers Ca^2+^ influx in the cytosol. IP_3_ can be synthesized by the Ca^2+^-dependent activity of Phospholipase C (PLC) *δ*. IP_3_ removal from the cytosol is described by a decay rate. IP_3_R dynamics is derived from the De Young & Keizer’s model [55]. Each IP_3_R has 3 binding sites: one to IP_3_ and two to Ca^2+^ (activating and inhibiting). The channel can thus be in 8 different states. The open state is {110}: IP_3_ and Ca^2+^ are bound to the activating sites and the Ca^2+^ inactivating site is unbound. Endogenous Ca^2+^ buffers are not explicitly modeled and their effect on Ca^2+^ dynamics is accounted for by decreasing the effective Ca^2+^ diffusion coefficient, as previously described [54, 56]. In a subset of simulations, GCaMPs6s, genetically-encoded Ca^2+^ indicators [29], were added to the cytosol and variations of [Ca-GCaMP] concentration, mimicking experimental Ca^2+^ imaging, were measured. For further details on the model assumptions, please refer to the original paper presenting the model [54]. We slightly altered this model to better describe and control IP_3_R-independent Ca^2+^ fluxes. To do so, IP_3_R-independent Ca^2+^ influx was modeled as an influx through Ca^2+^ channels at the plasma membrane, Ch_PM_. For simplicity, the number of Ch_PM_ channels was the same the total number of IP_3_R channels, *N*_IP3R_. Ca^2+^ influx rate at Ch_PM_ channels, *γ*, is 3 × 10^−2^ *s*^−1^.

The model was implemented using the STochastic Engine for Pathway Simulation (STEPS) (http://steps.sourceforge.net/) 3.5.0 [57, 58]. This software uses a spatialized version of Gillespie’s SSA algorithm [59] to perform exact stochastic simulations of reaction-diffusion systems. Simulations in STEPS allow the diffusion of molecules in 3D tetrahedral meshes and onto the surfaces of the mesh, such as the ER and plasma membrane. STEPS allows volume and surface reactions. Reactions can occur only between molecules within the same tetrahedron (volume reactions) or in adjacent triangle and tetrahedron (surface reactions). Boundary conditions were reflective. The simulation time was 100 s. The states and amounts of all molecular species were measured at each time step (1 ms).

#### 2.3.2 Neuronal stimulation simulation

Unless specified otherwise, glutamatergic transmission at the synapse was modeled and occurred at simulation time t=1 s. To do so, IP_3_ molecules were injected in tetrahedra below the plasma membrane of the PAP, emulating IP_3_ synthesis resulting from the activation of metabotropic glutamatergic receptors at the membrane of the PAP. Supplementary movie 4 presents a visualization of a simulation at neuronal stimulation time, in the d2s6a9b1 PAP mesh.

#### 2.3.3 Simulation code

Simulations were performed using the model of Ca^2+^ signals in fine processes from Denizot and collaborators [54], available at https://github.com/ModelDBRepository/247694.

The simulation code used in this study is available at https://github.com/adenizot/PAP-ER.

#### 2.3.4 Ca^2+^ residency time analysis

Ca^2+^ residency time was measured by performing n=16 simulations for each value of median d_ERPM_. In each simulation, for each IP_3_R, we injected a single Ca^2+^ ion in the IP_3_R site nanodomain. These regions of interest contained 15-16 tetrahedra each, consisting of the tetrahedron in contact with the IP_3_R triangle, neighboring tetrahedra, and tetrahedra in contact with the latter. Then, we tracked the time it took for the Ca^2+^ ion to diffuse away from the nanodomain.

#### 2.3.5 Ca^2+^ peak detection and characterization

Ca^2+^ peaks were considered initiated and terminated when Ca^2+^ concentration increased above and decreased below peak threshold, respectively. Peak threshold was [*Ca*]_b_ + *nσ*_Ca_, where [*Ca*]_b_ is the basal Ca^2+^ concentration and *σ*_Ca_ is the standard deviation of the [Ca^2+^] histogram in the absence of neuronal stimulation. n varied depending on the signal/noise ratio of the simulation of interest, notably when measuring Ca-GCaMP signals, noisier than free Ca^2+^ signals (see *e.g.* Fig 3C). Ca^2+^ peak frequency, duration, and amplitude were measured in each simulation. Ca^2+^ peak duration corresponds to the time between peak initiation and termination. Ca^2+^ peak amplitude corresponds to the maximum number of Ca^2+^ ions measured during peak duration. Peak amplitude is expressed as the number of Ca^2+^ ions (# Ca) in the cytosol of the whole PAP. Ca^2+^ peak frequency corresponds to the amount of peaks detected during the simulated time. The number of IP_3_R peak opening events was recorded at each time step, in the whole PAP.

### 2.4 Statistical analysis

Data analysis and statistics were performed using open-access and open-source software: the SciPy and Pandas Python libraries. Data visualization was performed using Seaborn and Matplotlib Python libraries. The sample size for each analysis, n, is described in the figure legend. Before statistical analysis, the normality of the data distribution was inferred using the Shapiro-Wilk test. The relationship between Ca^2+^ peak characteristics and parameter values was inferred using one-way ANOVA if values followed a Gaussian distribution, Kruskal-Wallis one-way ANOVA otherwise. The linear relationship between two datasets was evaluated using Spearman’s correlation coefficient. The test and p-value, p, associated with each analysis is described in the legend of the associated figure or in the main text.

## 3 Results

### 3.1 Quantification of the geometrical properties of hippocampal tripartite synapses

To characterize the geometrical properties of tripartite synapses, we used a 220 *µm*^3^ (7.07 *µ*m x 6.75 *µ*m x 4.75 *µ*m) hippocampal astrocytic volume from the CA1 stratum radiatum region reconstructed from a perfectly isotropic EM stack (6 nm voxel resolution) [51]. Elements from the neuropil, i.e. boutons, dendritic spine heads, and post-synaptic densities (PSDs), were also reconstructed. Following the workflow presented in Fig. 1A, forty-four excitatory and two inhibitory tripartite synapse meshes were reconstructed, containing all elements belonging to the astrocyte and to the neuropil within a cube of 1.5 *µ*m edge length (3.375 *µm*^3^) centered at the center of mass of the PSD (Supplementary Movie 1). Five of those tripartite synapse meshes are displayed in Fig. S1. Among these meshes, seventeen were located at the borders of the 220 *µm*^3^ astrocytic volume and thus could not be fully reconstructed. They were thus excluded from the quantitative analysis. The surface area, volume, and surface-volume ratio (SVR) of each synaptic element, i.e. the PAP, astrocytic ER, spine, and bouton, of the remaining twenty-seven fully reconstructed excitatory tripartite synapses are presented in Fig. 1C, D, F, respectively, and in Supplementary Table S1. Interestingly, we found a strong positive correlation between spine head and bouton volume (Fig. 1E), hallmark of synapse stability [60]. The large PAP SVR and smaller spine head and bouton SVRs reported here are in line with previous reports (see e.g. [49, 61] and Fig. S2). The distance between each vertex on the PAP membrane and the center of mass of the PSD was measured in each of the 27 meshes (Fig. 1B), providing a quantification of the distribution of the astrocyte around the synapse. Our results highlight the diverse distances between PSDs and PAPs belonging to a single cell. In line with previous studies [8, 62, 48], PAP membrane vertices could be as close as 5 nm to the PSD, with an average distance between the PSD and the closest PAP vertex of 65 nm. Importantly, PAPs were closer to the PSD than boutons and spines (Fig. 1G). The distance between PAPs and neuropil structures did not vary depending on the presence of ER in the PAP. Importantly, PAPs were closer to larger boutons and larger spines (Fig. 1H). Supplementary figures S3 and S4 display the distributions of the distances between each vertex of the membranes of PAPs and the closest spine (Fig. S3) or bouton (Fig. S4) vertex. Finally, we found that small PAP-PSD distances were correlated with large PAP surface area (Fig. 2H) and small bouton and spine surface areas (Fig. S5). Note that PAPs that were closer to boutons were also closer to spines and that bouton and spine size were positively correlated, while no correlation was found between PAP size and bouton or spine head size (Fig. S5). Overall, we report a high variability of the geometrical properties of PAPs belonging to the same astrocyte, which are correlated to the size of the neighboring synaptic elements.

**Figure 1:**
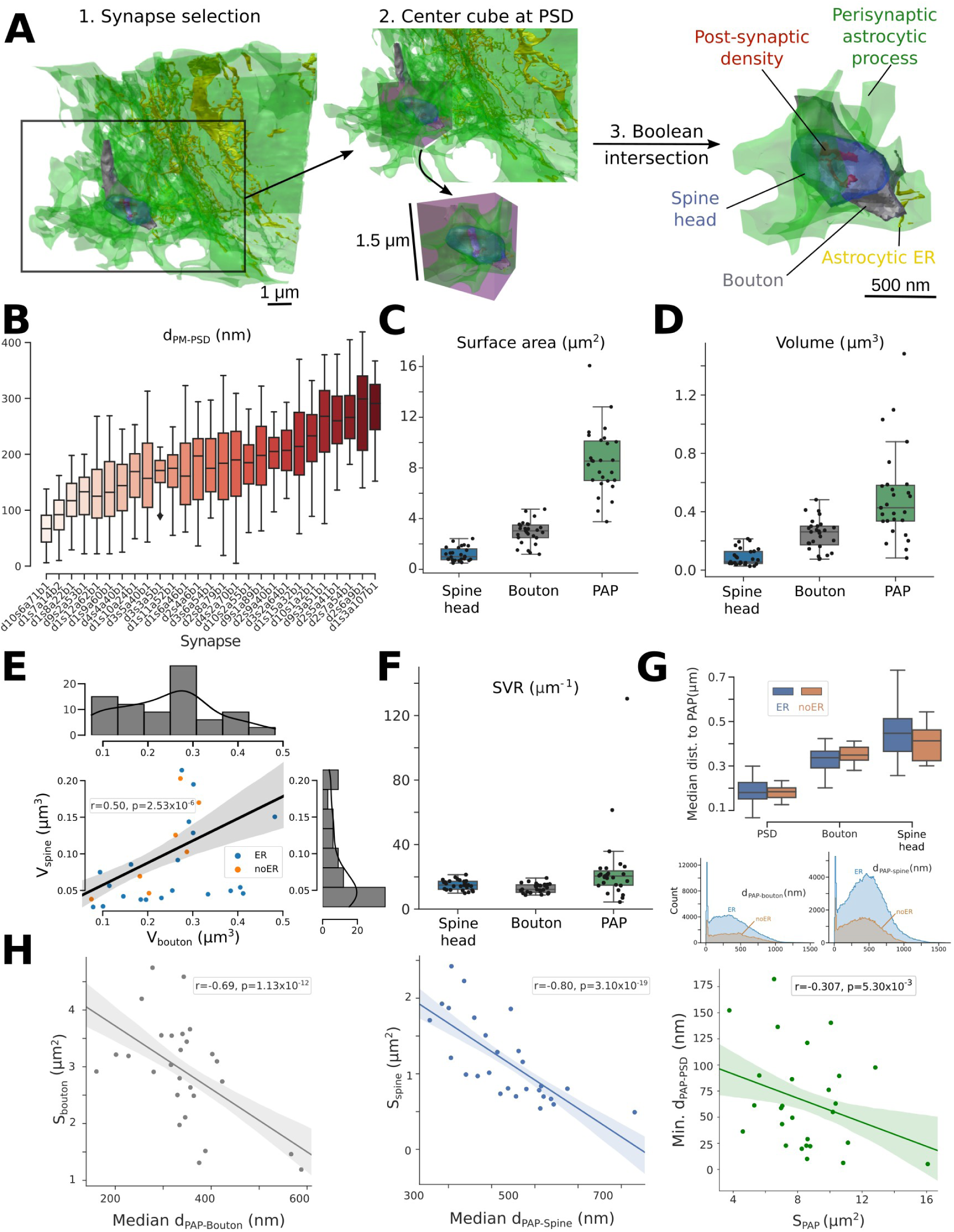
Quantification of the geometrical properties of hippocampal tripartite synapses. (A) Diagram presenting the workflow to extract tripartite synapse meshes (illustrated with synapse d10s1a2b1): 1. Synapses in contact with the 220 *µm*^3^ astrocytic volume were selected one by one. 2. A cube of 1.5 *µ*m edge length (3.375 *µm*^3^) was created and centered at the center of mass of the post-synaptic density (PSD, red). 3. Boolean intersection between the neuronal and astrocytic objects and the cube resulted in the isolation of the elements of the tripartite synapse mesh: the perisynaptic astrocytic process (PAP, green), the astrocytic endoplasmic reticulum (ER, yellow), the bouton (grey) and the spine (blue). This workflow resulted in the creation of 44 excitatory and 2 inhibitory tripartite synapse meshes. (B) Boxplots presenting the distribution of the minimum distance between each vertex on the PAP membrane and the center of mass of the PSD, measured in the 27 excitatory tripartite synapse meshes fully reconstructed in this study. (C-F) Boxplots presenting the distribution of spine, bouton, and PAP surface area (C), volume (D), and surface-volume ratio (F). (E) Scatterplot illustrating the increase of Spine volume with Bouton volume. Tripartite synapses in which PAPs contained some ER are represented with blue dots, orange otherwise. (G) Boxplots of median distances between PSD, Bouton, & spine heads to PAPs with (blue, n=20) or without (orange, n=7) ER (top). Distribution of PAP-bouton (left) & PAP-spine (right) distances differentiating PAPs with (blue, n=20) & without (orange, n=7) ER. (H) Scatterplots presenting the variation of median PAP-bouton distance with bouton surface area (left), median PAP-spine distance with spine surface area (middle), and minimum PAP-PSD distance with PAP surface area (right). Scatterplots are presented with a linear regression fit. Spearman correlation coefficients, r, and p-values, p, are displayed onto each regression plot, n=27.

### 3.2 Geometrical properties of the endoplasmic reticulum in perisynaptic astrocytic processes

Because of the nanoscopic size of most PAPs, the Ca^2+^ pathways that regulate astrocytic Ca^2+^ microdomain activity at tripartite synapses remain to be uncovered. Notably, the presence of ER in PAPs is controversial [48, 26, 50, 63]. We have thus investigated the presence and geometrical properties of the ER in the PAPs from the 27 fully reconstructed excitatory tripartite synapse meshes presented in Fig. 1.

75 % of PAPs contained some ER (Fig.2A-D), which challenges previous observations that reported that tripartite synapses are devoid of astrocytic ER. ER surface area, volume, and SVR were highly variable between PAPs of the same cell (Fig. 2C). Importantly, bouton, spine, and PAP surface area, volume, and SVR did not differ depending on the presence of ER in the PAP (Supplementary Fig. S6). In addition, we characterized the vicinity of the astrocytic ER to the synapse. To do so, we measured the distance between each vertex on the ER membrane to the center of mass of the PSD (n=20). We found that ER-PSD distance varies drastically from synapse to synapse (Fig. 2F-G) and can be as little as 70 nm, far below the *>* 0.5 *µ*m ER-PSD distance reported previously [48, 25]. The closest ER vertex was on average 432 nm away from the center of mass of the PSD. Interestingly, the larger the surface area of the ER, the closer it was to the PSD (Fig. 2G). Astrocytic ER was closer to the PSD in PAPs with higher surface area (Fig. 3G). ER surface area and ER-PSD distance were not correlated to the surface area of the PAP, spine, or bouton (Fig. S7A-B).

**Figure 2:**
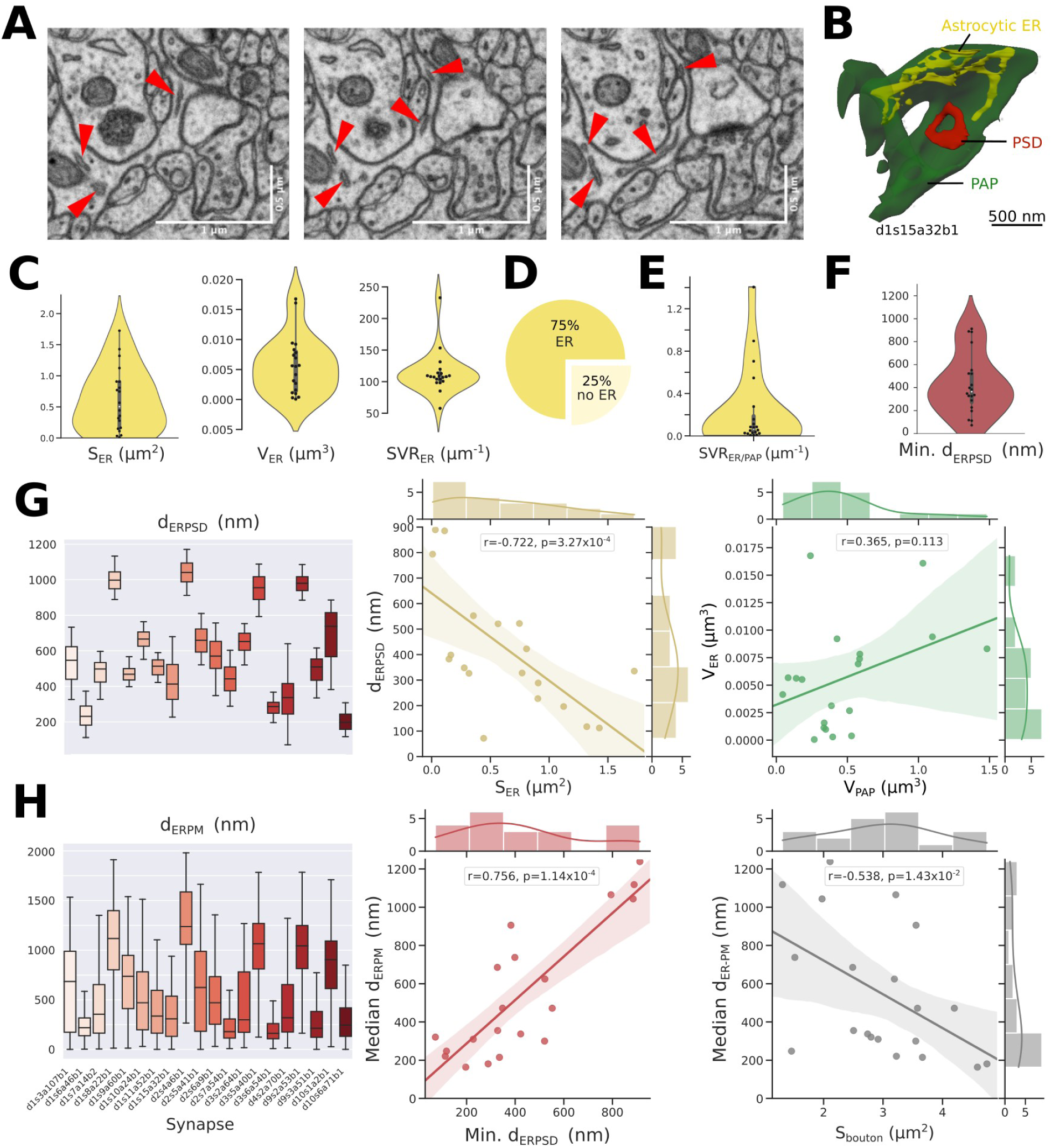
Characterization of the geometrical properties of the ER in PAPs. **(A)** Representative FIB-SEM images highlighting the presence of some endoplasmic reticulum (ER, ref arrowheads) in perisynaptic astrocytic processes (PAPs). (B) Image of the d1s15a32b1 PSD (red) and the neighboring PAP (green), that contains some ER (yellow). (C) Violin plots illustrating the distribution of ER surface area (left), volume (middle) and surface volume ratio (SVR, right) within PAPs, n=20. (D) Among the 27 fully reconstructed PAP meshes extracted, 75 % contained some ER. (E) Distribution of the ratio between the ER surface area and PAP volume (n=20). (F) Distribution of the minimum ER-PSD distance in PAPs, n=20. The lowest ER-PSD distance measured was 70 nm (synapse d4s2a70b1). (G) (Left) Boxplot presenting the distribution of the distance between each ER membrane vertex and the center of mass of the PSD in each PAP, d_ERPSD_, n=20. (Middle) Scatterplot presenting the negative correlation between the minimum ER-PSD distance and ER surface area. (Right) There is no strong correlation between PAP and ER volume. (H) (Left) Boxplot presenting the distribution of the distance between each PAP plasma membrane (PM) vertex and the closest ER vertex, d_ERPM_, n=20. Scatterplots presenting the variation of the median d_ERPM_ in PAPs as a function of the minimum d_ERPSD_ (Middle) and bouton surface area (Right). Scatterplots are presented with univariate kernel density estimation curves and a linear regression fit. Spearman correlation coefficients, r, and p-values, p, are displayed onto each regression plot, n=20.

We next aimed at quantifying ER-PM distance, d_ERPM_. To do so, we measured the distance between each vertex on the plasma membrane (PM) and the closest vertex on the ER. We found that d_ERPM_ is highly variable in PAPs from a single cell, with a median d_ERPM_ from around 200 nm to 1200 nm (Fig. 2H). Not surprisingly, median d_ERPM_ was negatively correlated with ER and PAP surface area (Fig. S7C). Interestingly, d_ERPM_ was negatively correlated to bouton surface area (Fig. 2H, p = 0.01). Importantly, we found that PAPs closer to the synapse were characterized by a lower median ER-PM distance (Fig. 2H, p = 1.14 *× 10*^−4^). Note that there was no correlation between d_ERPM_ and spine surface area (Fig. S7C).

Overall, our results highlight that, in the hippocampal CA1 stratum radiatum astrocyte volume reconstructed in this study, most astrocytic nanoscopic compartments that interact with synapses, PAPs, contained some ER. ER shape was highly variable, and was distributed closer to the plasma membrane in PAPs that were closer to the synapse. These correlational observations could have strong implications on ER-dependent Ca^2+^ signaling in PAPs resulting from synaptic transmission.

### 3.3 Reaction-diffusion simulations reveal different spatiotemporal properties of Ca^2+^ signals in PAPs of the same cell

PAPs are characterized by highly diverse sizes and distributions of the ER (Fig. 2), which could affect ER-mediated Ca^2+^ signals in PAPs. Because of their nanoscopic size, measuring Ca^2+^ activity and deciphering the involvement of ER-mediated signals in individual PAPs in live tissue is extremely challenging [28]. The mechanistic link between the geometrical properties of the ER and the spatiotemporal properties of Ca^2+^ microdomain signals in PAPs is thus unclear and hard to test experimentally. Yet, understanding the mechanisms that govern PAP activity is critical to deepen our understanding of neuron-astrocyte communication. Here, we use the PAP meshes extracted from EM presented in Fig. 2 together with a spatial stochastic model of Ca^2+^ signaling adapted from the model of Denizot and collaborators [54] to investigate the mechanistic link between ER shape and Ca^2+^ microdomain activity in PAPs. Ca^2+^ influx in the PAP cytosol in the model is mediated by Inositol 3-Phosphate (IP_3_) receptors on the membrane of the ER and by Ca^2+^ channels at the plasma membrane, Ch_PM_. The reactions modeled are presented in Fig. 3A, Supplementary Tables S2, S3, and in the Methods section. Neuronal activity was simulated at t=1 s by infusing 50 IP_3_ molecules at the PM of the PAP. The implementation of this model with STEPS software [64] allows simulations to be carried out in tetrahedral meshes in three spatial dimensions, such as the ones reconstructed in this study. Representative Ca-GCaMP traces, corresponding to the concentration of Ca^2+^ bound to Ca^2+^ indicators added to the cytosol of the model, display spatio-temporal characteristics similar to Ca^2+^ signals measured in organotypic hippocampal astrocytic cultures [65] (Fig. 3B).

**Figure 3:**
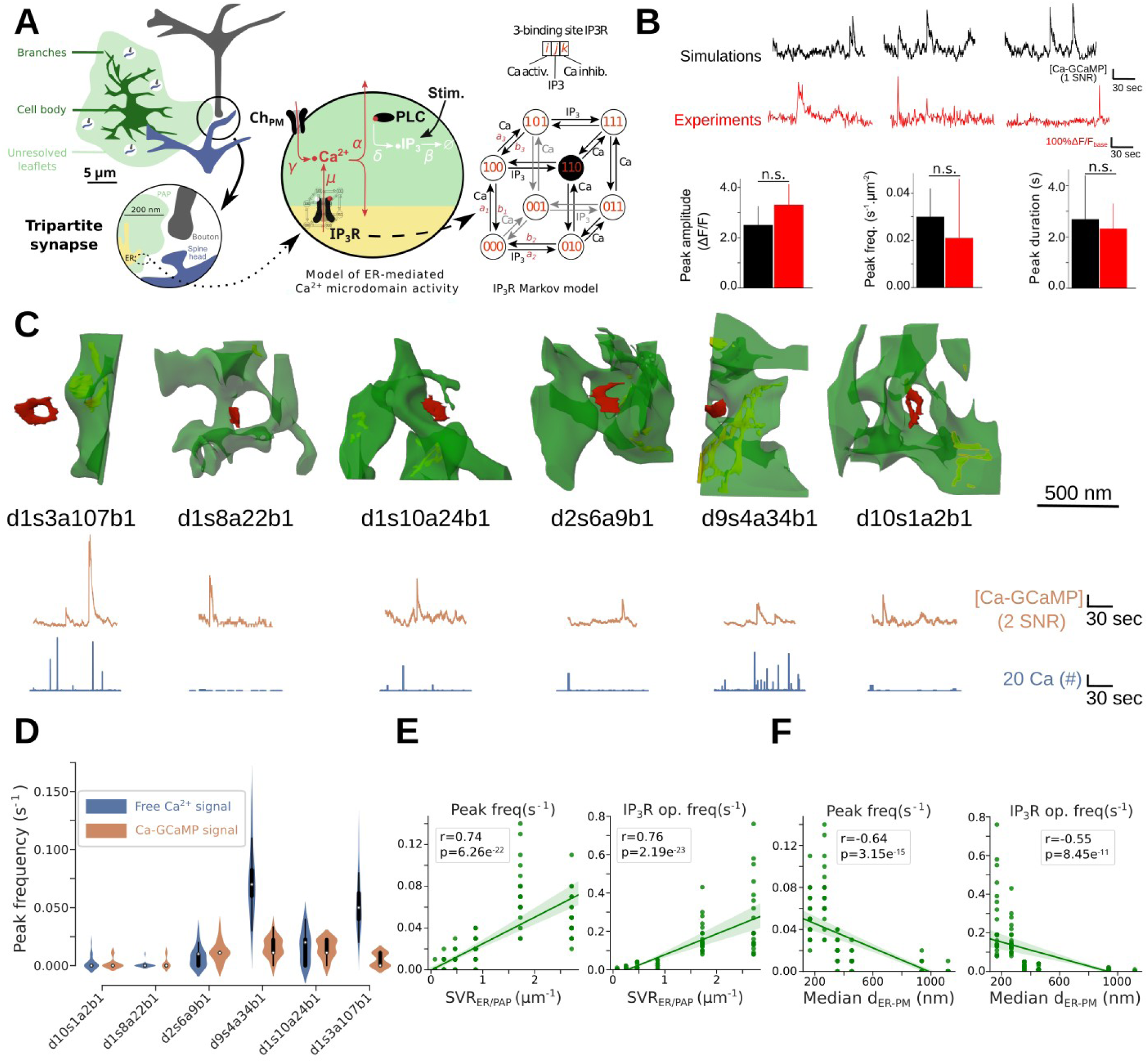
Reaction-diffusion simulations reveal different spatio-temporal properties of Ca^2+^ signals between PAPs of the same cell. (A) Schematic representation of the model of Ca^2+^ signaling in PAPs used in this study. The model is stochastic, spatially-extended and simulations can be performed in complex 3D meshes. Ca^2+^ influx into the cytosol results from Ca^2+^ channels on the plasma membrane and from IP_3_R channels on the ER. At t = 1 s, 50 IP_3_ molecules were injected at the plasma membrane of the PAP, simulating neuronal activity. The reactions modeled are detailed in Supplementary Table S2, parameter values and initial conditions in Table S3. (B) Representative Ca-GCaMP traces from simulations in a cylindrical mesh, 200 nm in diameter, 1 *µ*m long (top, black) and experiments (middle, red, [54]). SNR: signal-to-noise ratio (see Methods). (bottom) Ca^2+^ peak amplitude, frequency, and duration of simulated (black) and experimental signals (red) were similar. Data are presented as mean ± standard deviation, n= 20 and 29 for *in silico* and experimental traces, respectively. (C) Images of the 6 PAP meshes in which simulations were performed: d1s3a197b1, d1s8a22b1, d1s10a24b1, d2s6a9b1, d9s4a34b1, and d10s1a2b1 (top) and representative Ca-GCaMP (middle, orange) and free Ca^2+^ (bottom, blue) traces in each mesh. Free Ca^2+^ signals were measured in separate simulations, where no GCaMP was added into the cytosol of the PAP. IP_3_R channels and Ca^2+^ channels at the plasma membrane, Ch_PM_, were randomly distributed onto the ER membrane and plasma membrane, respectively. (D) Quantification of peak frequency of free Ca^2+^ (left, blue, n=20) and Ca-GCaMP (right, orange, n=20) signals measured *in silico* in 3D meshes of the PAPs presented in panel C. (E) Peak frequency (left) and IP_3_R opening frequency (right) are positively correlated with the ratio between the ER surface area and the cytosolic volume, SVR_ER*/*PAP_. (G) Peak frequency (left) and IP_3_R opening frequency (right) are negatively correlated with the median distance between each vertex on the PAP plasma membrane and the closest vertex on the ER membrane, *d*_ERPM_. Plots are presented with univariate kernel density estimation curves and a linear regression fit. Spearman correlation coefficient, r, and p-value, p, are displayed onto each regression plot.

We performed simulations in six PAP meshes reconstructed from electron microscopy, characterized by various geometrical properties of the ER: d1s3a107b1, d1s8a22b1, d1s10a24b1, d2s6a9b1, d9s4a34b1 and d10s1a2b1 (Fig. 3C, Table 1). To do so, meshes were pre-processed to allow their use in reaction-diffusion simulations. The pre-processing workflow is described in Fig. S8 and in the Methods section. Ca-GCaMP and free Ca^2+^ signals, in simulations with and without Ca^2+^ indicators in the cytosol, respectively, were measured in d1s3a107b1, d1s8a22b1, d1s10a24b1, d2s6a9b1, d9s4a34b1 and d10s1a2b1 PAP meshes. A simulation in PAP d9s4a34b1 is shown in Supplementary movie 2. Representative traces are displayed in Fig. 3C. Peak frequency (Fig. 3D), duration, and amplitude (Fig. S9) varied greatly depending on the mesh.

In accordance with previous studies [54, 66], Ca-GCaMP and free Ca^2+^ signals displayed different spatio-temporal properties (Fig. 3D and S9). These results suggest that the diverse geometrical features of PAPs and ER reported in Fig. 1 and 2, respectively, are correlated with different Ca^2+^ microdomain spatio-temporal properties. d1s3a107b1, d1s8a22b1, d1s10a24b1, d2s6a9b1, d9s4a34b1 and d10s1a2b1 PAPs displayed different ratios between the ER surface area and the cytosolic volume, SVR_ER*/*PAP;_, and median ER-PM distance d_ERPM_. Interestingly, Ca^2+^ peak frequency, IP_3_R opening frequency (Fig. 3 E-F), and Ca^2+^ peak amplitude and duration (Fig. S9C-D) were positively correlated with SVR_ER*/*PAP_ and negatively correlated with d_ERPM_. As the computational model explicitly describes the cytosolic compartment and the ER surface area but ignores intra-ER dynamics, we focused this study on the geometrical properties that alter ER surface properties. Consequently, geometrical features such as the ER volume or the fraction of PAP volume occupied by the ER, although positively correlated to Ca^2+^ peak activity (Fig. S10), were not considered in this study.

### 3.4 Ca^2+^ microdomain activity in PAPs increases with ER surface-PAP volume ratio

As Ca^2+^ peak frequency and IP_3_R opening frequency were positively correlated with the ratio between the ER surface area and the PAP volume, SVR_ER*/*PAP_ (Figure 3E), we next aimed at inferring the causal relationship between SVR_ER*/*PAP_ and Ca^2+^ microdomain activity. To do so, we created meshes with various ER surface area, while maintaining ER and PAP shapes. The original mesh was extracted from the 220 *µm*^3^ astrocytic volume, located at the vicinity of the d9s3a51b1 PSD and referred to as PAP1 (Fig. S11). Meshes with various SVR_ER*/*PAP_ were created from PAP1 by rescaling the ER using Blender software. Tetrahedral PAP meshes were then created following the mesh pre-processing workflow described in Fig. S8, resulting in the creation of PAP1_v_, PAP1_w_, PAP1_x_, PAP1_y_ and PAP1_z_ meshes (Fig. 4A). The geometrical properties of those meshes are presented in Table 1.

Spontaneous IP_3_R opening frequency (Fig. 4B), Ca^2+^ peak frequency (Fig. 4C), duration (Fig. S12A), and amplitude (Fig. S12B) increased with SVR_ER*/*PAP_. Interestingly, neuronal stimulation resulted in an increase of IP_3_R opening frequency (Fig. 4B) and was encoded in peak frequency (Fig. 4C) but did not significantly alter peak amplitude and duration (Fig. S12A-B). The increased IP_3_R opening and peak frequency with SVR_ER*/*PAP_ are not surprising as ER surface area S_ER_ increases with SVR_ER*/*PAP_ in those meshes. As IP_3_R density was constant across simulations, increasing S_ER_ thus resulted in an increase of the amount of IP_3_R channels with SVR_ER*/*PAP_. The total number of IP_3_R channels, *N*_IP3R_, thus was 24, 120, 240, 360 and 460, in PAP1_v_, PAP1_w_, PAP1_x_, PAP1_y_ and PAP1_z_ meshes, respectively. However, simulations with the same number of IP_3_R channels as PAP1_z_, 460, were performed in PAP1_w_, PAP1_x_, PAP1_y_, and PAP1_z_ meshes (Supplementary Fig. S13) and confirmed that SVR_ER*/*PAP_ influences IP_3_R opening frequency and thus Ca^2+^ peak frequency. Simulations in these meshes replicated the correlation between SVR_ER*/*PAP_ and Ca^2+^ peak frequency and IP_3_R opening frequency observed in Fig. 3. No correlation was found between SVR_ER*/*PAP_ and peak amplitude or duration (Fig. S13C-D). Note that no Ca^2+^ signals were detected in PAP1_v_ mesh.

**Figure 4:**
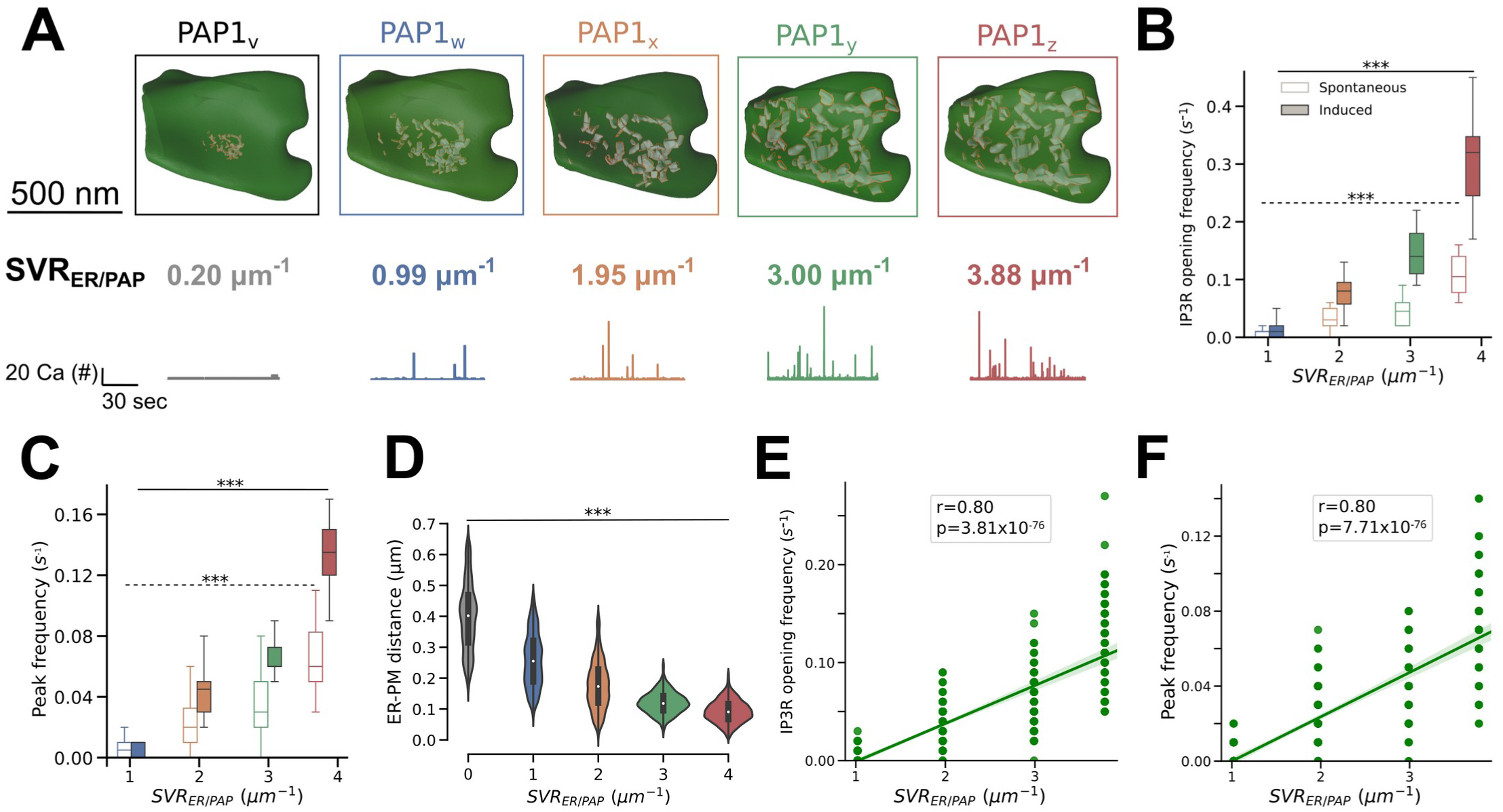
Ca^2+^ microdomain activity in PAPs increases with ER surface-PAP volume ratio. (A) (Top) Images of the different PAP meshes created to investigate the effect of the ratio between ER surface area and PAP volume, SVR_ER*/*PAP_, on Ca^2+^ microdomain activity: PAP1_v−z_. Meshes were obtained by rescaling the ER object in PAP1, located at the vicinity of the d9s3a51b1 PSD (Supplementary Fig. S11). Geometrical features of the meshes are presented in Table 1. (Bottom) Representative free Ca^2+^ traces measured in PAP1_v_ (grey), PAP1_w_ (blue), PAP1_x_ (orange), PAP1_y_ (green) and PAP1_z_ (red). IP_3_R opening frequency (B, ANOVA, p = 9.70 *× 10*^−83^ and p = 7.34 *× 10*^−65^ for spontaneous and induced Ca^2+^ signals, respectively) and Ca^2+^ peak frequency (C, ANOVA, p = 3.59 *× 10*^−80^ and p = 5.88 *× 10*^−128^ for spontaneous and induced Ca^2+^ signals, respectively) increase with SVR_ER*/*PAP_. (D) Quantification of the decrease of ER-PM distance *d*_ERPM_ with SVR_ER*/*PAP_ in PAP1_v−z_ meshes. IP_3_R opening frequency (E) and peak frequency (F) were positively correlated with SVR_ER*/*PAP_. Plots are presented with univariate kernel density estimation curves and a linear regression fit. Spearman correlation coefficient, r, and p-value, p, are displayed onto each regression plot.

Importantly, we noticed that increasing SVR_ER*/*PAP_ in PAP1 resulted in a decrease of the median distance between the ER and the plasma membrane (PM) in the PAP, *d*_ERPM_ (Fig. 4D), so that IP_3_R opening frequency and Ca^2+^ peak frequency were positively correlated with *d*_ERPM_ (Fig. 4E-F). Moreover, *d*_ERPM_ was also positively correlated with IP_3_R opening and Ca^2+^ peak frequency in meshes from Fig. 3. To differentiate the effect of *d*_ERPM_ from the effect of SVR_ER*/*PAP_ on Ca^2+^ dynamics, we next developed an algorithm to alter ER-plasma membrane distance independently of the shape, surface area, and volume of the ER and PAP.

### 3.5 Ca^2+^ microdomain activity is altered by the spatial distribution of the ER

To discern the effect of SVR_ER*/*PAP_ from the effect of d_ERPM_ on Ca^2+^ microdomain activity in PAPs reported in Fig. 3 and 4, we implemented an algorithm that generates realistic tetrahedral 3D meshes of PAPs characterized by various distributions of the ER within the same PAP with constant SVR_ER*/*PAP_. The workflow is presented in Fig. 5A. Briefly, the ER is split into small portions, then resized to match the total ER surface area of the original mesh. A simulation of n frames is then generated in Blender, which alters the location of the ER objects within the PAP. Each frame is thus characterized by a unique distribution of the ER objects within the PAP, while ER and PAP shape, surface area, volume, and SVR are constant across frames (Supplementary movie 3). The mesh processing workflow presented in Fig. S8 is then automatically applied to each frame of interest. This workflow allows the creation of numerous 3D PAP meshes characterized by various d_ERPM_, that can be used for reaction-diffusion simulations in 3D. The workflow successfully produced realistic tetrahedral PAP meshes characterized by various d_ERPM_, where *d*_ERPM_ decreased and the surface area of the ER at contact sites increased with frame number (Fig. 5B, workflow applied to PAP d1s15a32b1). The Blender file, python script, and parameter values used to generate these meshes are available at https://github.com/adenizot/PAP-ER.

**Figure 5:**
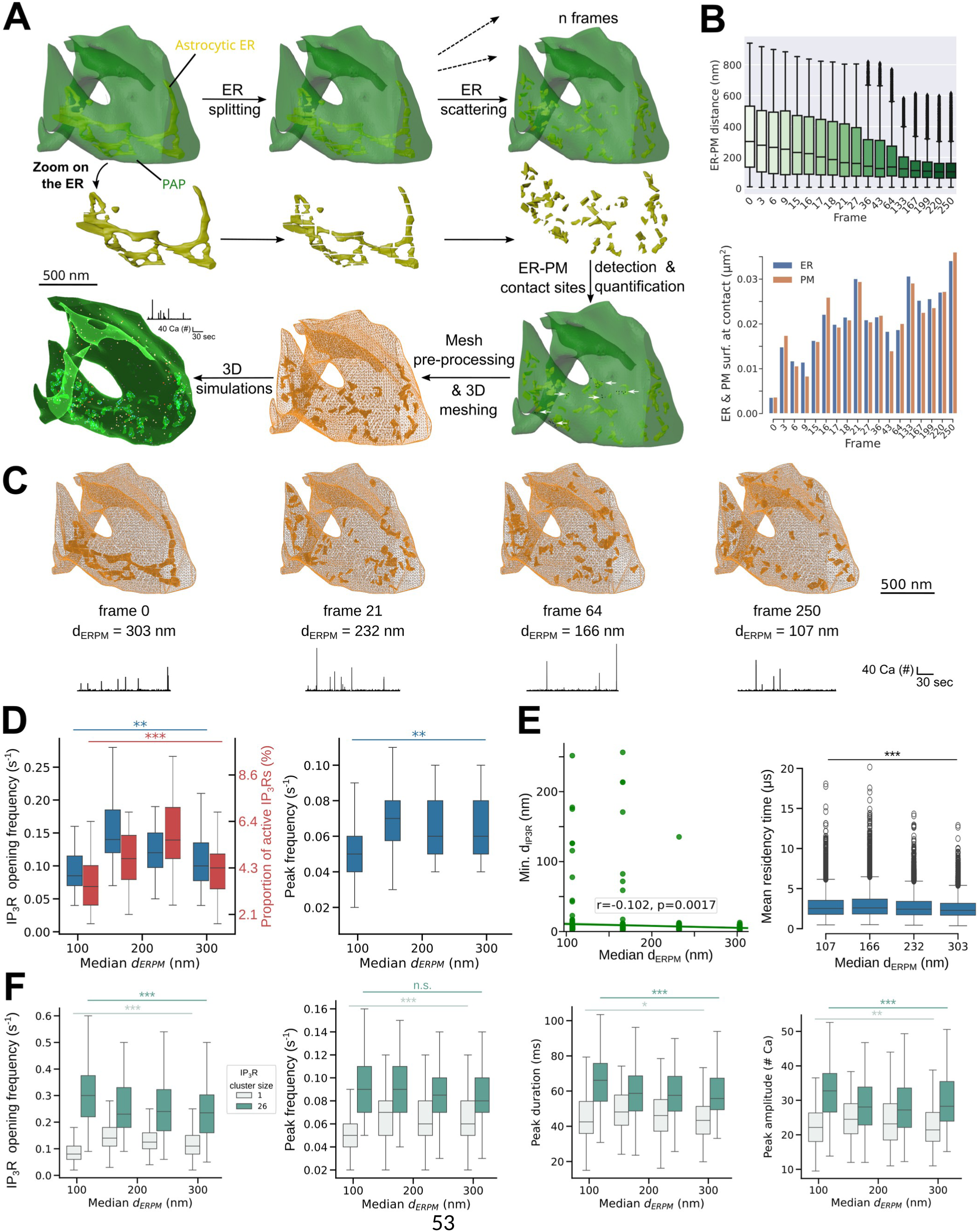
The spatial distribution of the ER dictates Ca^2+^ microdomain activity in perisynaptic astrocytic processes. (A) Schematic representing the workflow of the algorithm developed to create synthetic realistic PAP meshes in 3 spatial dimensions with various ER distributions and constant shape, volume and surface area of PAP and ER, used on the PAP mesh d1s15a32b1. The ER is split and a simulation with n frames is generated in Blender, in which ER objects are subject to physical forces that alter their spatial distribution. The n frames are thus characterized by different locations of the ER elements within the PAP, with constant ER and PAP shapes. The pipeline detects, quantifies and exports in a text file the distance between each vertex at the plasma membrane (PM) and the closest vertex at the membrane of the ER. A point cloud can be created to visualize the ER vertices at ER-PM contact sites (ER-PM distance *d*_ERPM_ ≤ 20 nm, white arrows). The mesh pre-processing workflow presented in Fig. 3C is then applied to the mesh of each desired frame. The resulting 3D tetrahedral meshes can then be used for 3D reaction-diffusion simulations. (B) (Top) Quantification of the distance between each PM vertex and the closest ER vertex in different frames of the simulation generated by the workflow presented in panel A. (Bottom) Quantification of the ER (blue, left) and PM (orange, right) surface area at ER-PM contact sites, in frames of the simulation generated by the workflow presented in panel A. (C) Images and representative free Ca^2+^ traces in different meshes created from PAP d1s15a32b1 using the automated workflow presented in panel A: d1s15a32b1_fr0_, d1s15a32b1_fr15_, d1s15a32b1_fr64_, and d1s15a32b1_fr250_, characterized by diverse ER distributions within the PAP with constant PAP and ER shapes, volumes and surface areas. (D) IP_3_R opening frequency (left, blue), the proportion of active IP_3_Rs (left, red), and Ca^2+^ peak frequency (right) increased with median *d*_ERPM_ (ANOVA, p = 0.0032, p = 8.19 *× 10*^−4^, and p = 0.024, respectively). (E) (left) The minimum distance between two adjacent IP_3_R sites decreased with median *d*_ERPM_. (right) Mean Ca^2+^ residency time in IP_3_R site nanodomains decreases as median *d*_ERPM_ increases. (F) (left) While in the absence of IP_3_R clusters IP_3_R opening frequency increased with the median d_ERPM_ (light green, ANOVA, p = 1.62 *× 10*^-13^, n=100), it was larger for smaller values of d_ERPM_ when ER-PM contact sites were equipped with IP_3_R clusters (dark green, p = 9.9*× 10*^-5^). (middle left) Peak frequency increased with d_ERPM_ in the absence of IP_3_R clusters (light green, p = 5.87 *× 10*^-9^) while no effect was observed upon IP_3_R clustering (dark green, p = 0.17). Similarly, Ca^2+^ peak duration (middle right) and amplitude (right) slightly increased with median d_ERPM_ in the absence of clusters (light green, p = 0.014 and p = 5.26 *× 10*^-3^ for peak duration and amplitude, respectively) while clustering of the Ca^2+^ channels resulted in increased peak duration and amplitude as the median d_ERPM_ decreased (dark green, p = 4.29 *× 10*^-4^ and p = 1.95 *× 10*^-4^, respectively).

Representative Ca^2+^ traces in meshes with various median *d*_ERPM_ are presented in Fig. 5C. IP_3_R channels were located at ER-PM contact sites, *i.e.* on the triangles of the ER surface that were the closest to a plasma membrane triangle. First, we checked that splitting the ER did not significantly alter Ca^2+^ activity in PAP meshes (Supplementary Fig. S14), which is possible because intra-ER Ca^2+^ dynamics is not considered in the model. Simulations in these meshes suggest that increasing median *d*_ERPM_ can trigger an increase in IP_3_R opening frequency and Ca^2+^ peak frequency, when IP_3_Rs are located at ER-PM contact sites (Fig. 5D). This increase is associated with an increase in the proportion of IP_3_Rs that get active at least once during simulation time (Fig. 5D). Note that Ca^2+^ peak amplitude and duration slightly increased with median *d*_ERPM_ (Fig. 5F). These results were confirmed by simulations in another realistic PAP mesh (Fig. S15). Interestingly, this suggests an opposite effect of *d*_ERPM_ on Ca^2+^ peak properties to that observed in Fig. 3F. This indicates that SVR_ER*/*PAP_ was probably the main determining factor of the variability of simulated Ca^2+^ peak frequency in the PAP meshes extracted from EM presented in Fig. 3. The observed correlation between *d*_ERPM_ and Ca^2+^ peak frequency most probably reflected an indirect effect of SVR_ER*/*PAP_, as SVR_ER*/*PAP_ and *d*_ERPM_ were negatively correlated (Spearman correlation coefficient r=-0.81, p = 8.45 *× 10*^−11^).

The simulation results presented in Fig. 5 suggest that a distribution of the ER further away from the plasma membrane can amplify peak frequency in the absence of IP_3_R clustering. This may seem counter-intuitive as ER-PM contact sites are often described as hubs of signal amplification. We hypothesize that, in the absence of IP_3_R clusters, increasing d_ERPM_ increases the probability of Ca^2+^ ions to diffuse away from contact sites, and thus increases the probability that it can reach a new IP_3_R channel to activate. To test this hypothesis, we measured the minimum distance between each pair of neighboring IP_3_Rs, *d*_IP3R_. We found that *d*_IP3R_ decreased and was less variable as median d_ERPM_ increased, confirming our intuition (Fig. 5E). We also confirmed that d_ERPM_ at IP_3_R sites increased with median d_ERPM_ (Fig. S16A), which could allow a faster diffusion of Ca^2+^ away from ER-PM contact sites to activate nearest IP3Rs. Moreover, the larger d_ERPM_ at the IP_3_R site, the larger the IP_3_R activity (expressed as the number of opening events per IP_3_R site, Fig. S16B). To better understand the mechanisms responsible for this increased activity at IP_3_R sites located further away from the plasma membrane, we performed simulations in which we infused a single Ca^2+^ ion at the IP_3_R site nanodomain and measured its residency time (see Methods for details on the protocol). We observed that mean Ca^2+^ residency time decreased as d_ERPM_ increased, confirming our hypothesis that increasing d_ERPM_ increases the probability of Ca^2+^ ions to diffuse away from contact sites. Importantly, distributing IP_3_Rs in clusters at ER-PM contact sites led to an increase of IP_3_R activity and of Ca^2+^ peak properties as median *d*_ERPM_ decreased (Fig. 5F).

Overall, our results suggest that IP_3_R sites close to the plasma membrane are characterized by a restricted signal diffusion to neighboring IP_3_R sites, which can reduce or boost Ca^2+^ activity depending on the absence or presence of IP3R clusters, respectively. Our results thus nuance the view of ER-PM contact sites as local amplifiers of Ca^2+^ activity, conditioning this effect to the presence of clusters of Ca^2+^ channels.

## 4 Discussion

Here, we reconstructed 3D meshes of tripartite synapses from an isotropic high-resolution 220 *µm*^3^ hippocampal stratum radiatum EM dataset [51]. Quantitative analysis of those meshes highlighted the diverse geometrical properties of PAPs within a single astrocyte and revealed, contrary to a widespread assumption that PAPs are devoid of ER [48, 25, 49], that 75 % of PAPs contained some ER in this dataset. We found that PAPs were closer to the PSD when bouton surface area was low, which could result from the spatial constraints imposed by larger boutons, preventing the PAP from getting in close contact to the PSD. We observed that PAPs were closer to larger boutons and larger spines. Bouton volume is correlated with the number of pre-synaptic vesicles [67] and with the size of the active zone and the latter scales with release probability [68]. Spine head volume also correlates with the number of presynaptic vesicles anchored [69]. Thus, our data suggest that PAPs might be closer to spines and boutons of more active synapses. Our data further suggest that the ER is distributed closer to the plasma membrane in PAPs connected to larger boutons and thus more active synapses, which could, according to our simulations, impact Ca^2+^ microdomain signaling in the PAP and thus neuron-astrocyte communication. Future studies on the astrocyte nano-architecture will be important to test whether these features, observed in a single astrocyte of the CA1 region, differ with intra- and inter-regional astrocyte heterogeneity, in different physiological conditions, and in different species.

Reaction-diffusion simulations in the realistic PAP 3D meshes reconstructed in this study provided key insights into the effect of the diverse shapes and distributions of the ER observed in PAPs on microdomain Ca^2+^ activity. Notably, we reported that larger SVR_ER*/*PAP_ triggered increases in IP_3_R opening frequency and Ca^2+^ peak frequency. This could be due to intracellular diffusional barriers resulting from larger intracellular compartments, similarly to diffusional barriers mediated by the spongiform morphology of PAPs as reported previously [70]. Moreover, our simulations suggest that a distribution of the ER further away from the plasma membrane can amplify peak frequency in the absence of IP_3_R clustering. Our results suggest that this effect results from an increased probability of Ca^2+^ ions released at an open IP_3_R to reach a new IP_3_R channel. This is due to a decreased distance between adjacent IP_3_Rs and a decreased Ca^2+^ residency time when the ER is further away from the plasma membrane. Importantly, we show that the effect of the spatial distribution of the ER on Ca^2+^ dynamics strongly depends on the spatial distribution of IP_3_Rs, in particular their organization into clusters at ER-PM contact sites, as reported in HelA cells [71]. Future work studying the spatial distribution of IP_3_Rs in PAPs will thus be crucial to better understand the mechanisms regulating Ca^2+^ activity at tripartite synapses.

As reactive astrocytes, hallmark of brain diseases [72], are characterized by a remodelling of ER volume and shape [73], our results suggest that such geometrical remodeling of the ER could contribute to the altered astrocytic Ca^2+^ activity reported in pathological conditions [74].

Combining our detailed biophysical model of Ca^2+^ signals in PAPs, the PAP meshes that we reconstructed from EM, and the PAP meshes with various ER distributions produced by our automated mesh generator allowed us to fine-tune the spatial distribution of Ca^2+^ channels, monitor IP_3_R opening events at individual channels, while independently manipulating ER shape and distribution, providing new insights into some mechanisms governing Ca^2+^ signaling in PAPs. Notably, we predict how the ratio between ER surface area and PAP volume, and the spatial distribution of the ER shape Ca^2+^ microdomain signals at tripartite synapses. This study is the first to our knowledge to model Ca^2+^ activity in astrocytes within realistic shapes in 3D at the nanoscale that accounts for the complex and diverse spatial characteristics of Ca^2+^ stores in PAPs and shows how small differences in the 3D intracellular landscape such as the spatial distribution of the ER, can shape local signaling. Historically, modeling studies on PAPs, including our own, have been conducted in 1D, 2D, or in simple 3D shapes, such as cylinders [54, 70, 75, 76, 77, 78]. The 3D meshes provided by this study and the algorithm generating 3D PAP meshes with various ER distributions with constant shape and size pave the way for future modeling studies to investigate the mechanisms governing neuron-astrocyte communication at tripartite synapses.

The ultrastructural data presented in this study were reconstructed from 3D FIB-SEM electron microscopy, whose tissue processing protocol, in particular the chemical fixation, might not preserve adequately the extracellular space volume, and might alter the shape of fine astrocytic structures [79]. Importantly, this technique provides a static view of the ultrastructural properties of the tissue as it cannot be used to study live cells. However, EM provides the highest spatial resolution (6 nm isotropic here) to determine PAP and ER shape to date. The exact ultrastructure of the astrocytic ER in live tissue and the physiological relevance of the ER discontinuities sometimes observed in 3D reconstructions from EM in small sub-cellular compartments, such as the astrocytic ER of this dataset, are still unclear. *In vivo* and *in vitro* super-resolution studies have recently revealed that neuronal ER is continuous in healthy conditions [80, 81, 82] and undergoes rapid fission under cerebral ischemic conditions [83] or preceding excitotoxic cell death [84]. Recently, studies have shown that this apparent continuity results from a balance between fast fission and fusion events of the ER in dendritic spines *in vitro* [85] as well as *in vivo*, following somatosensory stimulation [81]. Interestingly, cortical spreading depolarization, which causes migraine aura, also triggers widespread ER fission *in vivo* [81]. Those results highlight the plasticity of the ER shape in neurons in live tissue. Whether such fission events occur in astrocytes and their potential contribution to astrocyte function remains to be uncovered. By generating 3D meshes with various ER splitting and scattering characteristics, the algorithm developed in this study could be used to replicate different scenarios of ER fission and investigate their effect on cellular activity under (patho-)physiological conditions, although here we have simply used it as a tool to infer the causal link between the distribution of the ER and the spatio-temporal properties of Ca^2+^ signals independently from other spatial properties.

The model used in this study describes in details the kinetics of ER-mediated Ca^2+^ signals while simplifying other Ca^2+^ sources and channels, such as mitochondria, the Na^+^/Ca^2+^ exchanger, transient receptor potential ankyrin 1 channels and L-type voltage gated channels [29, 47]. The model is therefore well-suited to study the effect of ER shape and distribution on Ca^2+^ activity but does not allow to study other astrocytic Ca^2+^ pathways. According to our predictions, the spatial distribution of Ca^2+^ channels can alter the spatio-temporal properties of Ca^2+^ microdomain signals in PAPs. Additional quantification of the Ca^2+^ channels expressed in PAPs, their density, location in live tissue, and the remodeling of these properties under (patho-)physiological conditions will thus be essential to better understand neuron-astrocyte communication at synapses. The recent advances in super-resolution techniques, notably single-particle tracking methods, provide a promising avenue to overcome current limitations in obtaining such data [23, 86]. Another approximation of the model is that endogenous buffers are not explicitly described. The effect of Ca^2+^ buffering on cytosolic Ca^2+^ dynamics is well documented [87]. Notably, their binding to Ca^2+^ ions results in a decreased diffusion distance of free Ca^2+^ ions [88]. Thus, for simplicity, this model describes the effect of buffers with a decreased effective diffusion coefficient of free Ca^2+^ ions. In the original study presenting the model, we showed that there was no significant difference between simulations with fast Ca^2+^ diffusion and slow explicit buffers and the model with an effectively reduced Ca^2+^ diffusion coefficient (Fig. S1 of Denizot et al. [54]). In accordance with previous reports [54, 89], our results highlight how Ca^2+^ indicators such as GCaMP, act as Ca^2+^ buffers and influence Ca^2+^ dynamics, notably increasing signal duration and decreasing peak frequency and amplitude. The local buffer concentrations dictate the decay of Ca^2+^ peaks. Buffers differ in their affinity, binding/unbinding to Ca^2+^ rates, and diffusional properties. Future research on astrocyte buffering capacity and buffer expression at the sub-cellular scale in various (patho-)physiological conditions will thus be essential to fully understand the regulation of astrocytic Ca^2+^ dynamics and homeostasis by Ca^2+^ buffers.

Here, we used a FIB-SEM dataset in the hippocampal CA1 stratum radiatum to analyze the 3D structure of tripartite synapses and of the ER inside perisynaptic astrocytic processes. It is important to note that the high resolution of the dataset (6 nm isotropic voxel size) allowed to reconstruct only a portion of a single astrocyte. Moreover, astrocyte morphology is highly complex, nanoscopic, dynamic, and diverse [90], so that the results presented in this study might not be generalizable. Recent pioneering vEM studies have started to unravel the organizing principles of astrocyte structure [63, 90, 91, 92]. Future research will be instrumental to characterize the ultrastructural properties of multipartite synapses in various species, brain regions, developmental stages, and pathological conditions, to determine the fundamental structural principles of astrocytes at the nanoscale. Such data will be crucial to increase our understanding of the now well-established astrocyte inter- and intra-cellular heterogeneity [93, 94, 95, 96] in health and disease. Acquiring and analyzing such high-resolution datasets is challenging and will require interdisciplinary efforts to develop novel computational and computer vision tools tailored to astrocytes to facilitate their segmentation, structural analysis, and data storage [97, 98]. Combining these insights into astrocyte structure with functional studies, for example using volume correlative light-electron microscopy [99], will contribute to increasing our understanding of the roles of astrocyte morphology.

Overall, this study provides new insights into astrocyte function at tripartite synapses by characterizing the shape and distribution of the ER in PAPs of an astrocyte in the hippocampal CA1 stratum radiatum and by shedding light into the mechanistic link between those features and microdomain Ca^2+^ activity at tripartite synapses. Notably, most PAPs in this dataset contained some ER. *In silico* experiments showed that the ER surface to PAP volume ratio, the spatial distribution of the ER within PAPs, and IP_3_R clustering finely regulate Ca^2+^ microdomain activity. The realistic 3D meshes of tripartite synapses provided in this study pave the way for new modeling studies of neuron-astrocyte communication in the synaptic micro-environment, allowing the study of various processes, such as glutamate spillover or gliotransmission. Such studies will be crucial to decipher whether the various nano-architectures displayed by tripartite synapses reflect distinct functional identities.

## Data availability statement

The data and code that support the findings of this study are openly available at https://zenodo.org/records/17106549 and https://github.com/adenizot/PAP-ER, respectively.

## Acknowledgments

This work used the computing resources of the Scientific Computing and Data Analysis section from the Okinawa Institute of Science and Technology. We thank Iain Hepburn and Weiliang Chen (OIST, Japan) for discussion and advice on STEPS software. We thank Pierre Magistretti (KAUST, Thuwal) for the financial support of MFVC and of CC, and Graham Knott for kindly sharing the original EM dataset. This work was supported by a JSPS (Japan Society for the Promotion of Science) Standard Postdoctoral Fellowship for Research in Japan (21F21733).

