## Supplemental materials for "The ultrastructural properties of the endoplasmic reticulum govern microdomain signaling in perisynaptic astrocytic processes"

### Supplementary materials

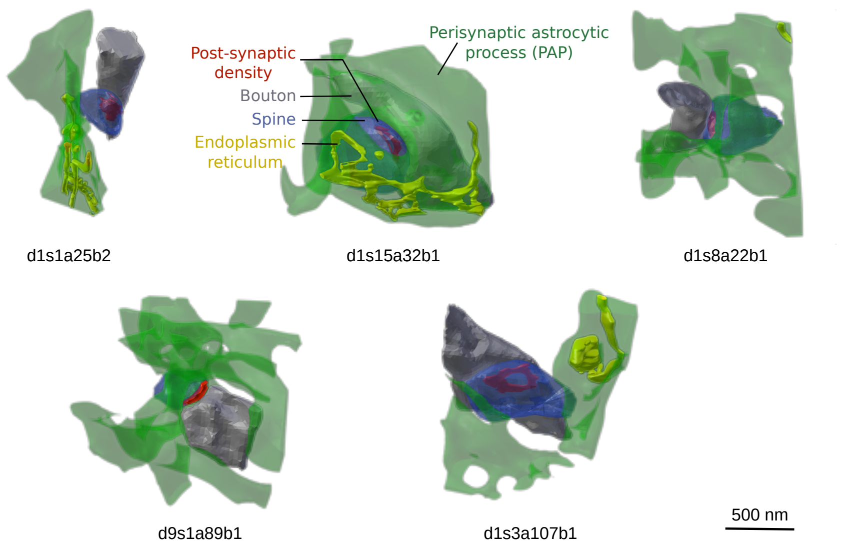


Fig S1: **Images of five representative 3D tripartite synapse meshes.** Screenshots of synapses d1s1a25b2, d1s15a32b1, d1s8a22b1, d9s1a89b1, d1s3a107b1, revealing their diverse geometrical properties. Perisynaptic astrocytic processes (PAPs) are in green, astrocytic ER in yellow, boutons in grey, spine heads in blue, and PSDs in red.

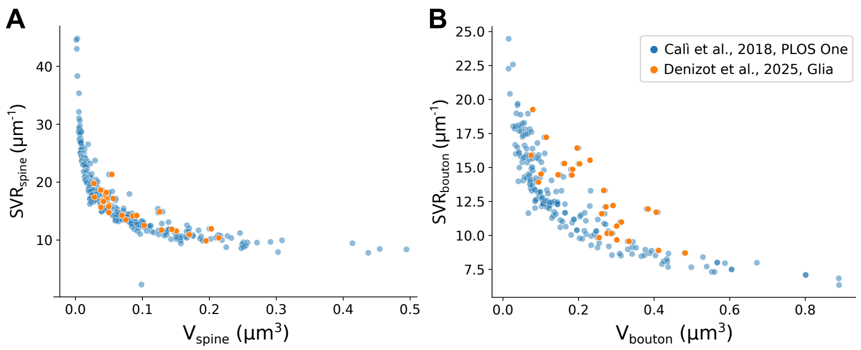


Fig S2: **Spine head and bouton geometrical properties are consistent with previous reports.** Spine head (A) and bouton (B) volume and surface-volume ratio (SVR) measured in this study (orange) are in line with measurements from Calì et al., 2018, PLOS One [1] (blue).

Table S1: **Characteristics of the twenty seven tripartite synapse meshes reconstructed from electron microscopy.** $V_{Bouton}$, $V_{Spine}$, $V_{PAP}$ and $V_{ER}$ are the bouton, spine, PAP and astrocytic ER volumes, respectively. $S_{Bouton}$, $S_{Spine}$, $S_{PAP}$ and $S_{ER}$ are the bouton, spine, PAP and astrocytic ER surface area, respectively.

| Synapse | $V_{bouton}$ ($\mu m^3$) | $V_{spine}$ ($\mu m^3$) | $V_{PAP}$ ($\mu m^3$) | $V_{ER}$ ($\mu m^3$) | $S_{bouton}$ ($\mu m^2$) | $S_{spine}$ ($\mu m^2$) | $S_{PAP}$ ($\mu m^2$) | $S_{ER}$ ($\mu m^2$) |
|---|---|---|---|---|---|---|---|---|
| d1s3a107b1 | 0.16 | 0.09 | 0.18 | 0.0056 | 2.49 | 1.21 | 3.76 | 0.32 |
| d1s6a46b1 | 0.20 | 0.05 | 0.24 | 0.017 | 3.22 | 0.81 | 5.31 | 1.43 |
| d1s7a14b2 | 0.25 | 0.05 | 0.59 | 0.0078 | 2.50 | 0.74 | 8.58 | 0.77 |
| d1s8a22b1 | 0.09 | 0.07 | 0.40 | 0.0003 | 1.31 | 1.01 | 8.61 | 0.03 |
| d1s9a60b1 | 0.08 | 0.03 | 0.34 | 0.0016 | 1.52 | 0.54 | 8.54 | 0.16 |
| d1s10a24b1 | 0.23 | 0.04 | 0.39 | 0.0031 | 3.58 | 0.67 | 7.67 | 0.36 |
| d1s11a52b1 | 0.18 | 0.04 | 0.59 | 0.0074 | 2.74 | 0.70 | 10.19 | 0.81 |
| d1s12a62b1 | 0.20 | 0.05 | 0.34 | - | 3.10 | 0.80 | 7.27 | - |
| d1s15a32b1 | 0.30 | 0.19 | 0.43 | 0.0092 | 2.92 | 1.92 | 6.98 | 0.90 |
| d2s4a6b1 | 0.15 | 0.04 | 0.53 | 0.0004 | 2.11 | 0.70 | 4.6 | 0.04 |
| d2s5a41b1 | 0.33 | 0.04 | 0.58 | 0.0069 | 3.19 | 0.80 | 6.76 | 0.75 |
| d2s6a9b1 | 0.48 | 0.15 | 0.52 | 0.0027 | 4.20 | 1.74 | 10.06 | 0.28 |
| d2s7a54b1 | 0.41 | 0.05 | 1.48 | 0.048 | 4.75 | 1.06 | 6.54 | 0.91 |
| d2s8a79b1 | 0.31 | 0.17 | 0.38 | - | 3.44 | 1.86 | 8.26 | - |
| d2s9a40b1 | 0.26 | 0.13 | 0.88 | - | 3.03 | 1.87 | 12.82 | - |
| d3s2a64b1 | 0.27 | 0.09 | 0.14 | 0.0056 | 3.55 | 1.30 | 8.60 | 0.57 |
| d3s3a5b1 | 0.18 | 0.07 | 0.20 | - | 2.64 | 0.99 | 5.62 | - |
| d3s5a40b1 | 0.30 | 0.13 | 0.27 | 5.4e-5 | 3.21 | 1.51 | 7.035 | 0.008 |
| d3s6a54b1 | 0.38 | 0.05 | 1.098 | 0.0094 | 4.59 | 0.78 | 7.65 | 1.02 |
| d4s2a70b1 | 0.28 | 0.21 | 0.045 | 0.0041 | 2.80 | 2.23 | 10.83 | 0.44 |
| d4s4a40b1 | 0.29 | 0.10 | 0.74 | - | 2.91 | 1.29 | 11.02 | - |
| d9s1a89b1 | 0.07 | 0.04 | 0.51 | - | 1.09 | 0.60 | 10.36 | - |
| d9s2a53b1 | 0.11 | 0.06 | 0.35 | 0.0010 | 1.97 | 0.97 | 8.78 | 0.11 |
| d9s3a51b1 | 0.41 | 0.05 | 1.032 | 0.016 | 3.66 | 0.84 | 9.93 | 1.72 |
| d10s1a2b1 | 0.27 | 0.20 | 0.55 | 0.0011 | 3.30 | 2.42 | 10.58 | 0.15 |
| d10s2a55b1 | 0.29 | 0.14 | 0.34 | - | 3.55 | 1.71 | 7.045 | - |
| d10s6a71b1 | 0.10 | 0.0017 | 0.083 | 0.0057 | 1.46 | 0.49 | 10.83 | 0.67 |

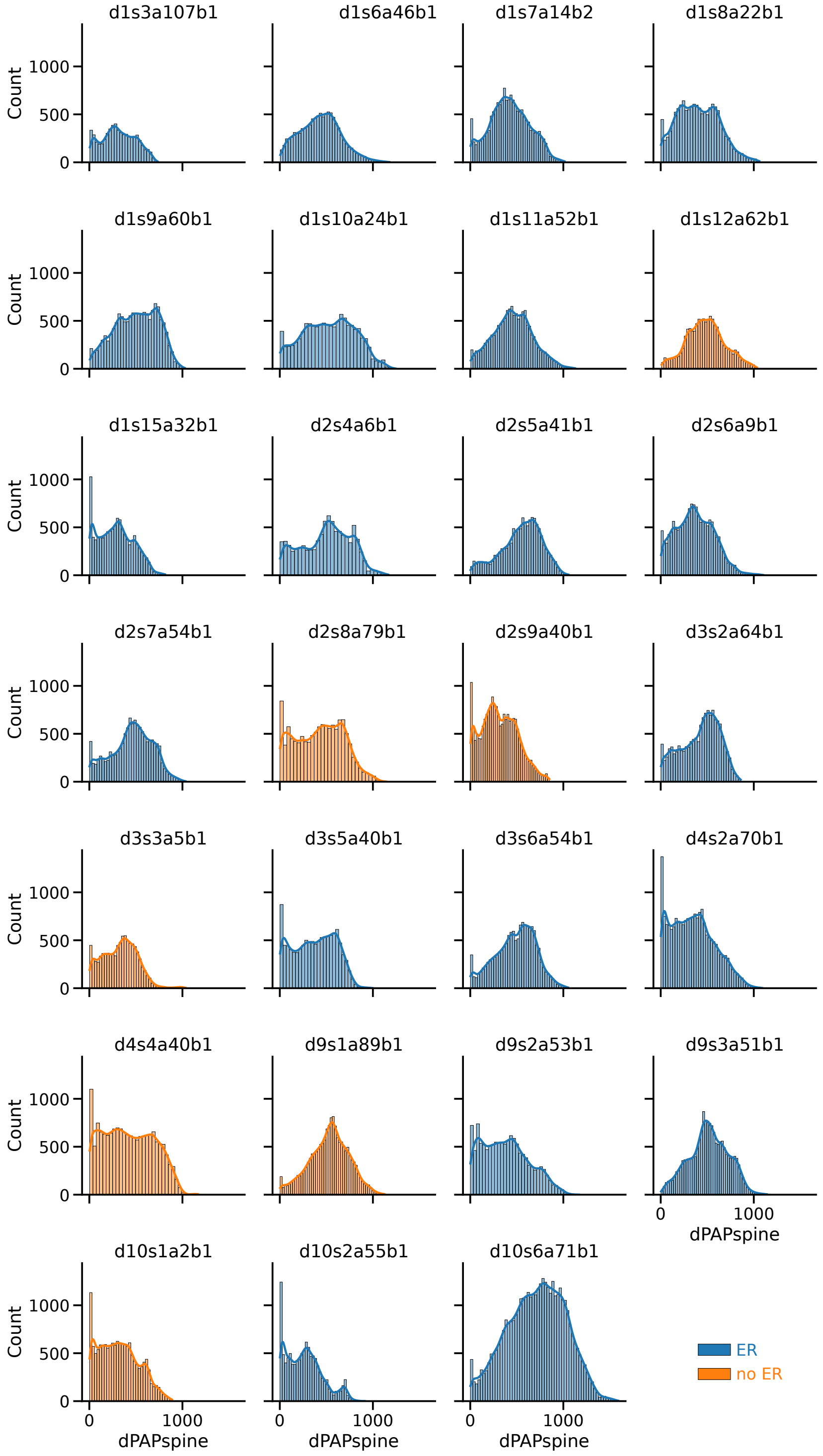


Fig S3: **Distribution of PAP-spine distances in individual synapses.** Distribution of the distances between triangles at the membrane of PAPs & spines in each of the 28 tripartite synapses analyzed in this study. Synapses in which PAPs contained some ER are represented in blue, orange otherwise. Histograms are presented with univariate kernel density estimation curves.

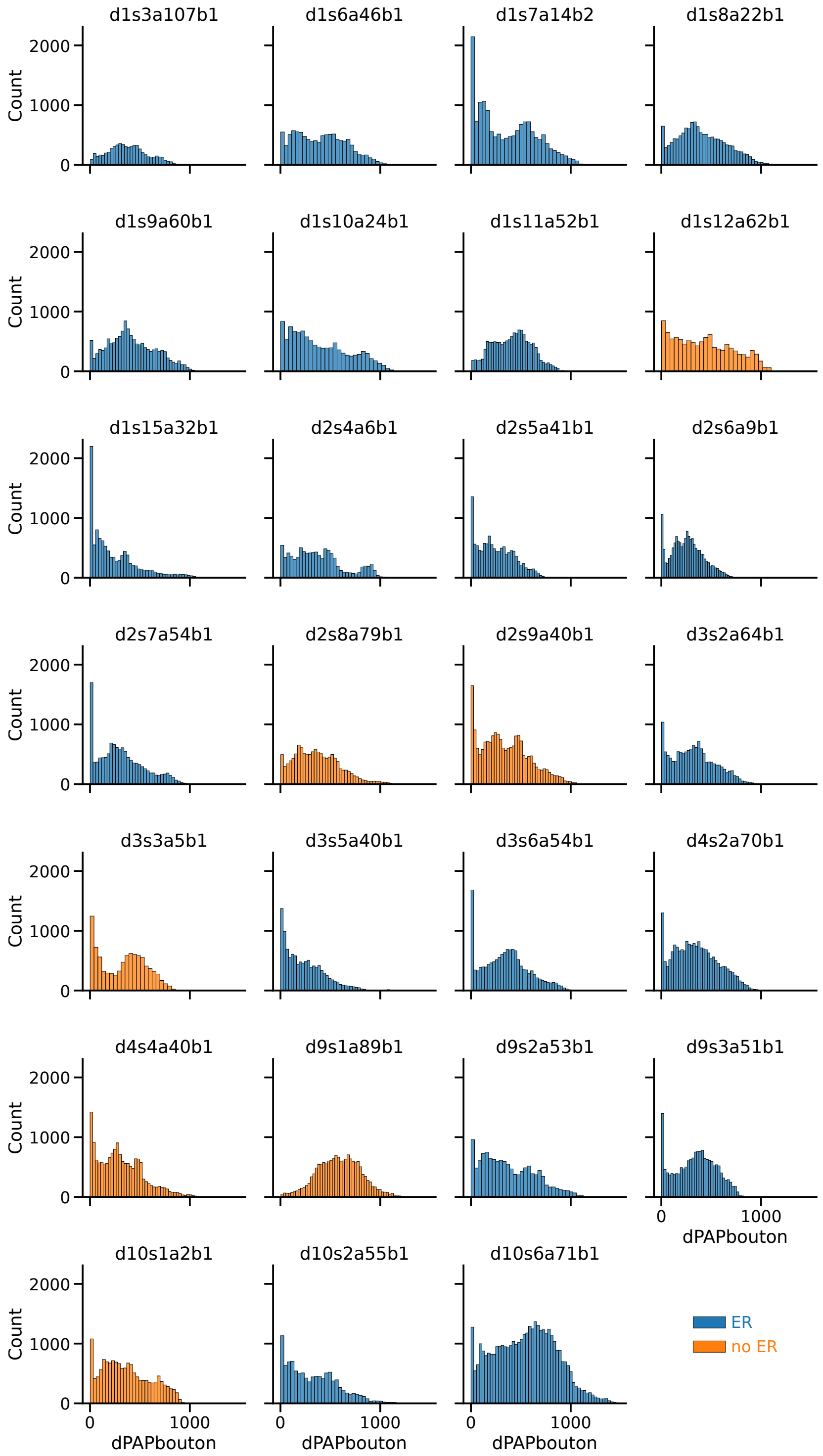


Fig S4: **Distribution of PAP-bouton distances in individual synapses.** Distribution of the distances between triangles at the membrane of PAPs & boutons in each of the 28 tripartite synapses analyzed in this study. Synapses in which PAPs contained some ER are represented in blue, orange otherwise. Histograms are presented with univariate kernel density estimation curves.

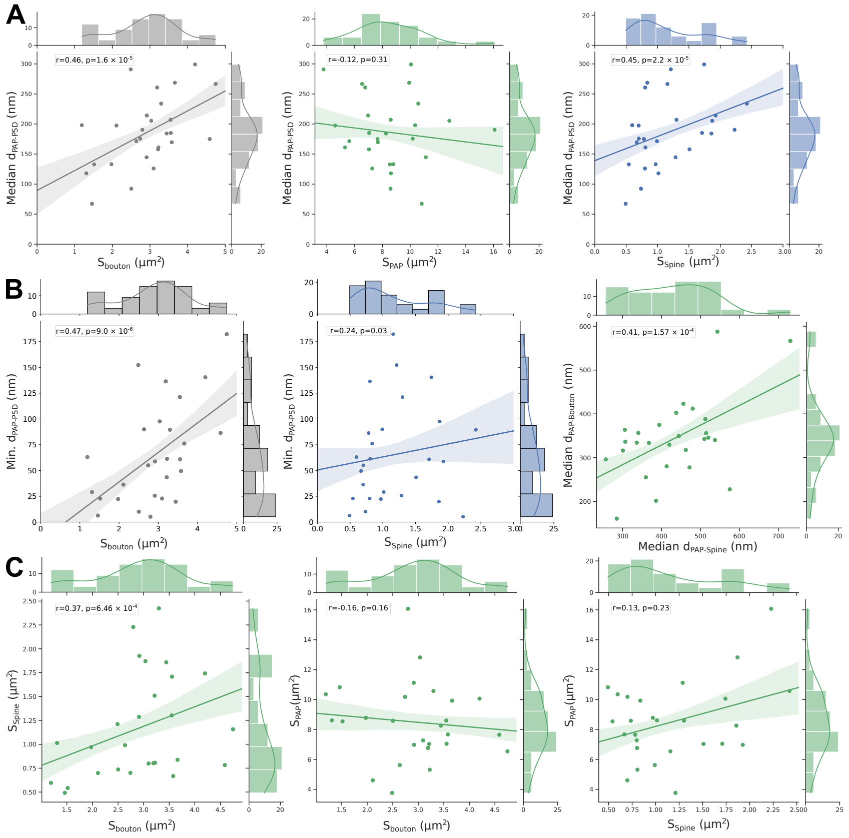


Fig S5: **PAP & synapse geometrical properties are correlated.** (A) Bouton (left) & spine (right) surface area increase with median PAP-PSD distance while PAP surface area decreases (center). (B) Minimum PAP-PSD distance increases as Bouton (left) & Spine (center) surface area increase. Median PAP distances to boutons and spines are positively correlated (right). (C) PAP & Bouton surface area are positively correlated to Spine surface area (left, right, respectively) while PAP surface area is negatively correlated to Bouton surface area (center). Plots are presented with univariate kernel density estimation curves and a linear regression fit. Spearman correlation coefficient, r, and p-value, p, are displayed onto each regression plot.

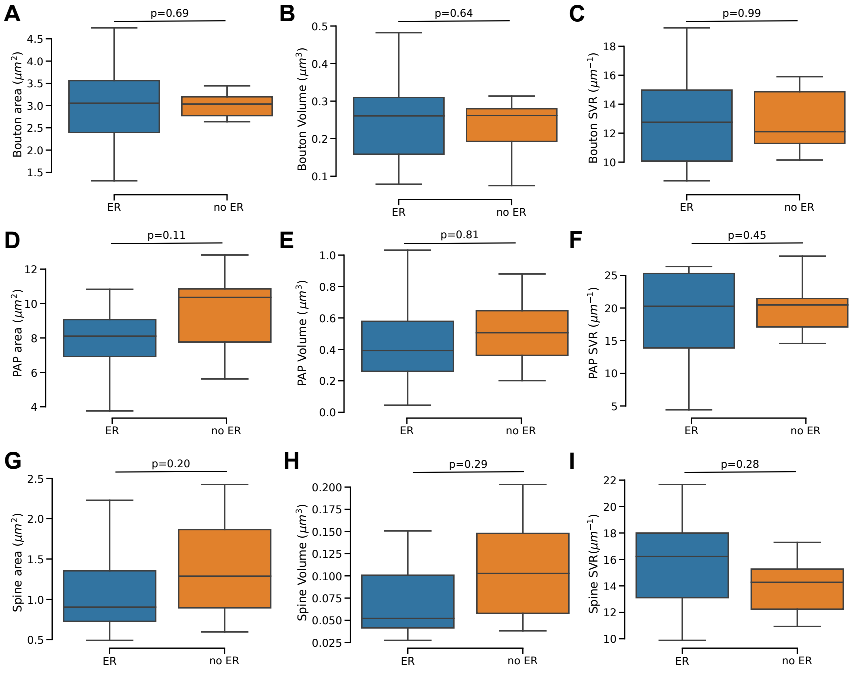


Fig S6: **Tripartite synapses with and without astrocytic ER display similar geometrical properties.** (A-C) Bouton surface area (A), volume (B) and SVR (C), PAP surface area (D), volume (E) and SVR (F), spine surface area (G), volume (H) and SVR (I) are not significantly different in synapses contacted by a PAP with ('ER', blue) vs without ER ('no ER', orange). p-values p (unpaired Student T-test), are displayed onto each plot. 'ER': n=20, 'noER': n=7.

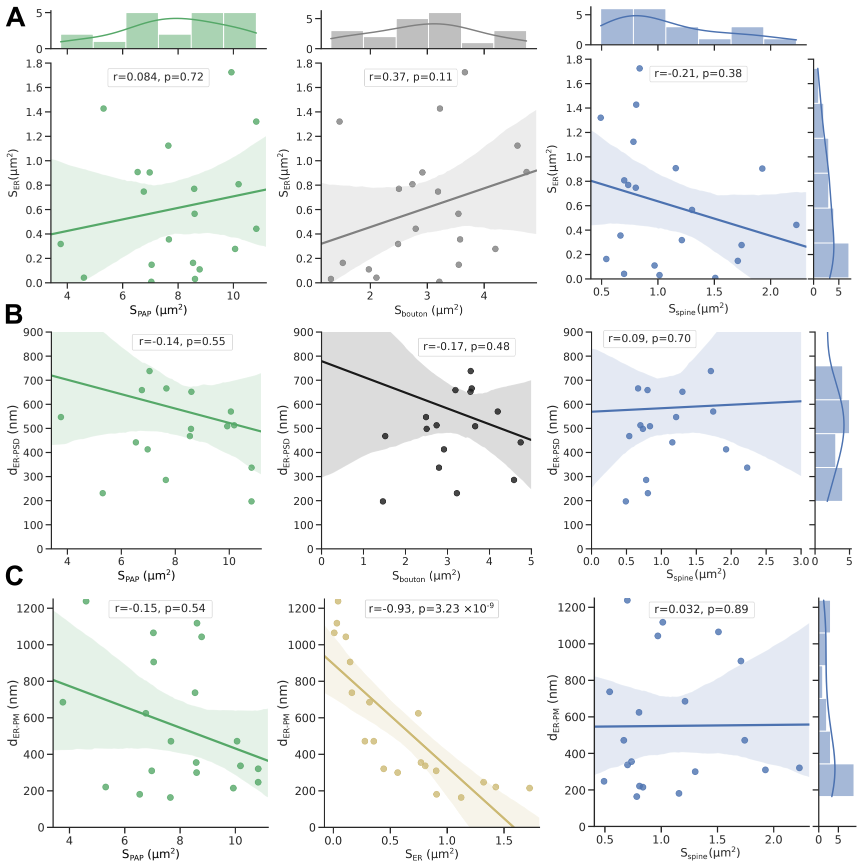


Fig S7: **Analysis of correlations between astrocytic ER & tripartite synapse geometrical properties.** (A) ER surface area $S_{ER}$ is not correlated to perisynaptic astrocytic process (PAP, left, green), bouton (middle, grey), and spine (right, blue) surface area. (B) The median distance between the ER and the center of mass of the post-synaptic density, $d_{ER-PSD}$, is not correlated to PAP (left, green), bouton (middle, grey), and spine (right, blue) surface area. (C) The median distance between each vertex on the PAP plasma membrane (PM) and the closest ER vertex, $d_{ER-PM}$, is not correlated to PAP (left, green), and spine (right, blue) surface area. As expected, $d_{ER-PM}$ decreases as $S_{ER}$ increases. Plots are presented with univariate kernel density estimation curves and a linear regression fit. Spearman correlation coefficient, r, and p-value, p, are displayed onto each regression plot.

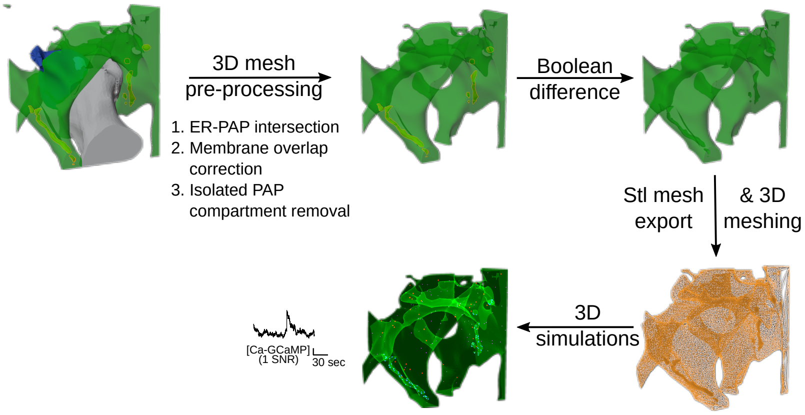


Fig S8: **Workflow to prepare the PAP meshes for 3D simulations illustrated on d2s6a9b1 mesh.** The geometrical features of the resulting PAP meshes are presented in Table 1 (see Methods section for details).

Table S2: **Reactions modeled.**

| **Reaction** | **Parameter** |
|---|---|
| $IP_3$ dynamics | |
| $IP_3 \to \emptyset$ | $\beta$ |
| $PLC\delta + Ca \to PLC\delta + Ca + IP_3$ | $\delta$ |
| $Ca^{2+}$ dynamics | |
| $Ca \to \emptyset$ | $\alpha$ |
| $Ch_{PM} \to Ch_{PM} + Ca$ | $\gamma$ |
| GCaMP6s | |
| GcaMP + Ca → GCaMP-Ca | $g_f$ |
| GcaMP + Ca ← GCaMP-Ca | $g_b$ |
| $IP_3R$ dynamics | |
| [000] → [100] | $a_1$ |
| [000] ← [100] | $b_1$ |
| [000] → [010] | $a_2$ |
| [000] ← [010] | $b_2$ |
| [000] → [001] | $a_3$ |
| [000] ← [001] | $b_3$ |
| [011] → [111] | $a_1$ |
| [011] ← [111] | $b_1$ |
| [001] → [011] | $a_2$ |
| [001] ← [011] | $b_2$ |
| [010] → [011] | $a_3$ |
| [010] ← [011] | $b_3$ |
| [110] → [111] | $a_3$ |
| [110] ← [111] | $b_3$ |
| [010] → [110] | $a_1$ |
| [010] ← [110] | $b_1$ |
| [101] → [111] | $a_2$ |
| [101] ← [111] | $b_2$ |
| [001] → [101] | $a_1$ |
| [001] ← [101] | $b_1$ |
| [100] → [110] | $a_2$ |
| [100] ←[110] | $b_2$ |
| [100] → [101] | $a_3$ |
| [100] ← [101] | $b_3$ |
| [110] → [110] + Ca | $\mu$ |

Table S3: **Parameter values and initial conditions.** The parameter values and initial conditions of the model are derived from the "GCaMP" and "No-GCaMP" models of [2, 3]. The model kinetic scheme is described in Figure 3A. Plasma membrane $Ca^{2+}$ channels, $Ch_{PM}$, were modeled as described in [3].

| **Parameter** | **Description** | **Value** |
|---|---|---|
| | $IP_3$ dynamics | |
| $IP_0$ | Initial $IP_3$ concentration | 120 nM |
| $D_{IP3}$ | $IP_3$ diffusion coefficient | 280 $\mu m^2.s^{-1}$ |
| $d_{plc}$ | PLC$\delta$ density at the plasma membrane | 2.463 $\times 10^3 \mu m^{-2}$ |
| $\delta$ | PLC$\delta$ max rate | 1 $s^{-1}$ |
| $\beta$ | $IP_3$ decay rate | $1.2 \times 10^{-4}$ $s^{-1}$ |
| | $Ca^{2+}$ dynamics | |
| $Ca_0$ | Initial $Ca^{2+}$ concentration | 120 nM |
| $D_{Ca}$ | $Ca^{2+}$ diffusion coefficient | 13 $\mu m^2.s^{-1}$ |
| $N_{ch}$ | Number of $Ch_{PM}$ channels | $N_{IP_3R}$ |
| $\gamma$ | $Ca^{2+}$ flux through $Ch_{PM}$ channels | $3 \times 10^{-2}$ $s^{-1}$ |
| $\mu$ | $Ca^{2+}$ flux through open $IP_3R$ | $6 \times 10^3$ $s^{-1}$ |
| $\alpha$ | $Ca^{2+}$ decay rate | 30 $s^{-1}$ |
| | $IP_3R$ | |
| $d_{IP_3R}$ | $IP_3R$ density on the ER membrane | 3.4 $\times 10^2 \mu m^{-2}$ |
| | $IP_3R$ binding | |
| $a_1$ | Activating Ca | $1.2 \times 10^6$ $M^{-1}.s^{-1}$ |
| $a_2$ | $IP_3$ | $4.1 \times 10^7$ $M^{-1}.s^{-1}$ |
| $a_3$ | Inhibiting Ca | $1.6 \times 10^4$ $M^{-1}.s^{-1}$ |
| | $IP_3R$ dissociation | |
| $b_1$ | Activating Ca | 50 $s^{-1}$ |
| $b_2$ | $IP_3$ | 400 $s^{-1}$ |
| $b_3$ | Inhibiting Ca | 100 $s^{-1}$ |
| | GCaMP6s (when applicable) | |
| $C_{GCaMP}$ | GCaMP6s concentration | 10 $\mu$M |
| $D_{GCaMP}$ | GCaMP6s diffusion coefficient | 50 $\mu m^2.s^{-1}$ |
| $g_f$ | GCaMP6s Ca binding rate | $7.78 \times 10^6$ $M^{-1}.s^{-1}$ |
| $g_b$ | GCaMP6s-Ca dissociation rate | 1.12 $s^{-1}$ |

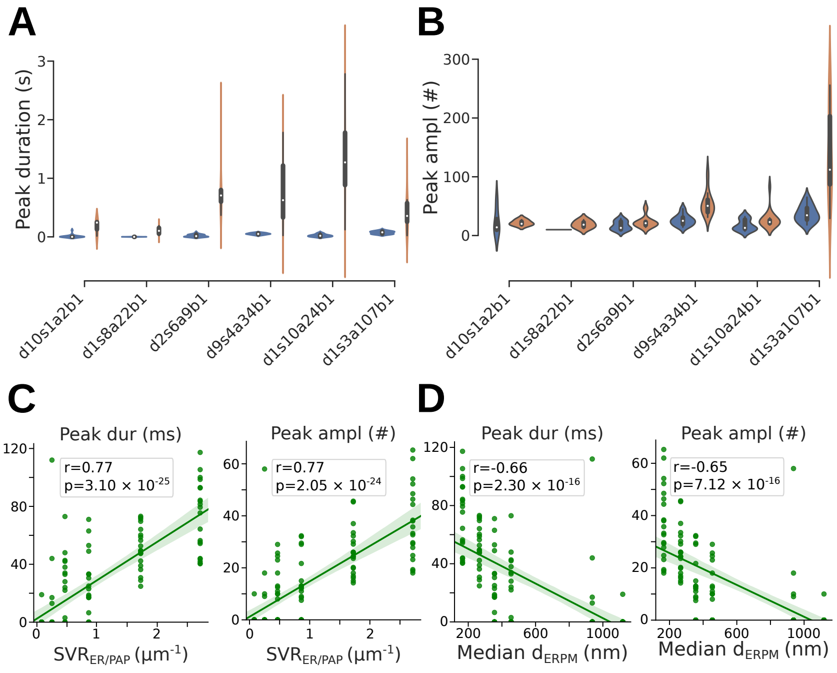


Fig S9: $Ca^{2+}$ **peak amplitude and duration vary between PAPs of the same cell.** Quantification of peak duration (A) and amplitude (B) of free $Ca^{2+}$ (left, blue, n=20) and Ca-GCaMP (right, orange, n=20) signals measured *in silico* in 3D meshes of the PAPs presented in Fig. 3C. (C) Peak duration (left) and amplitude (right) are positively correlated with the ratio between the ER surface area and the cytosolic volume, $SVR_{ER/PAP}$. (D) Peak duration (left) and amplitude (right) are negatively correlated with the median distance between each vertex on the PAP plasma membrane and the closest vertex on the ER membrane, $d_{ERPM}$. Plots are presented with univariate kernel density estimation curves and a linear regression fit. Spearman correlation coefficient, r, and p-value, p, are displayed onto each regression plot.

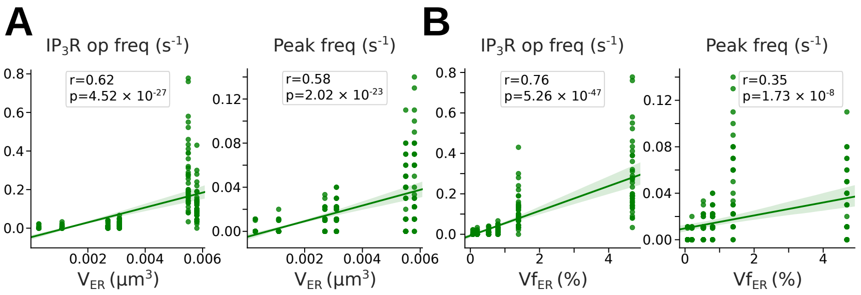


Fig S10: $Ca^{2+}$ **activity is correlated with ER volume and the fraction of PAP volume occupied by the ER.** (A) $IP_3R$ opening frequency (left) and $Ca^{2+}$ peak frequency (right) are positively correlated with the total ER volume $V_{\mathrm{ER}}$. (B) $IP_3R$ opening frequency (left) and $Ca^{2+}$ peak frequency (right) are positively correlated with the fraction of the cytosolic volume occupied by the ER, $Vf_{\mathrm{ER}}$. Plots are presented with univariate kernel density estimation curves and a linear regression fit. Spearman correlation coefficient, r, and p-value, p, are displayed onto each regression plot.

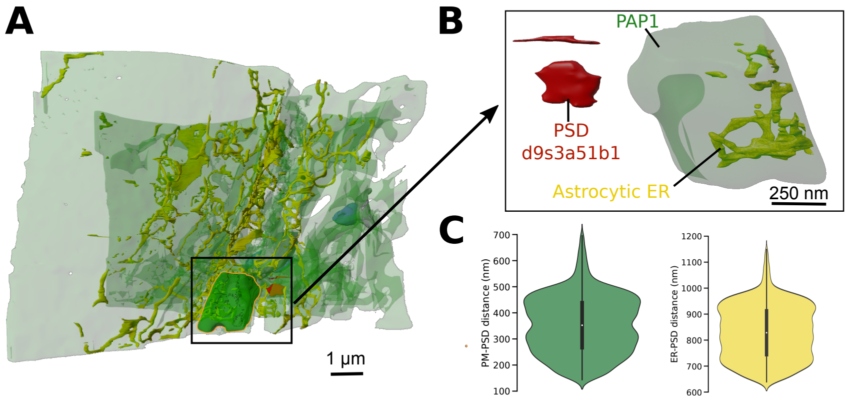


Fig S11: **Geometrical properties of PAP1 mesh.** (A) Visualization of the location of PAP1 within the 220 $\mu m^3$ hippocampal astrocytic volume. (B) Zoom on PAP1, with its neighboring PSD: d9s3a51b1. The ER was selected in Blender and rescaled. The resulting meshes, characterized by various $\mathrm{SVR}_{\mathrm{ER/PAP}}$ and constant PAP shape, were processed using the workflow presented in Fig. S8, resulting in the creation of the $\mathrm{PAP1}_{\mathrm{v-z}}$ meshes presented in Fig. 4. (C, left) Distribution of the distances between vertices on the plasma membrane (PM, green) of PAP1 and the center of mass of the d9s3a51b1 PSD. (C, right) Distribution of the distances between astrocytic ER vertices in PAP1 (yellow) and the center of mass of the d9s3a51b1 PSD.

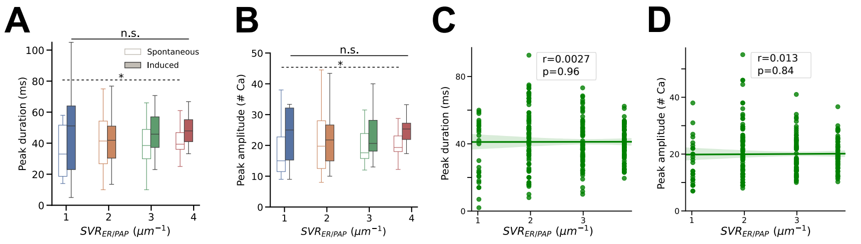


Fig S12: **Impact of the ER surface-volume ratio on** $Ca^{2+}$ **peak amplitude and duration in PAPs.** Spontaneous $Ca^{2+}$ peak duration (A) and amplitude (B) slightly increased with $SVR_{ER/PAP}$ (ANOVA, $p = 0.033$ and $0.035$ for duration and amplitude, respectively), while this effect was absent for signals induced by $IP_3$ infusion. Peak duration (C) and amplitude (D) are not correlated with $SVR_{ER/PAP}$. Plots are presented with univariate kernel density estimation curves and a linear regression fit. Spearman correlation coefficient, r, and p-value, p, are displayed onto each regression plot.

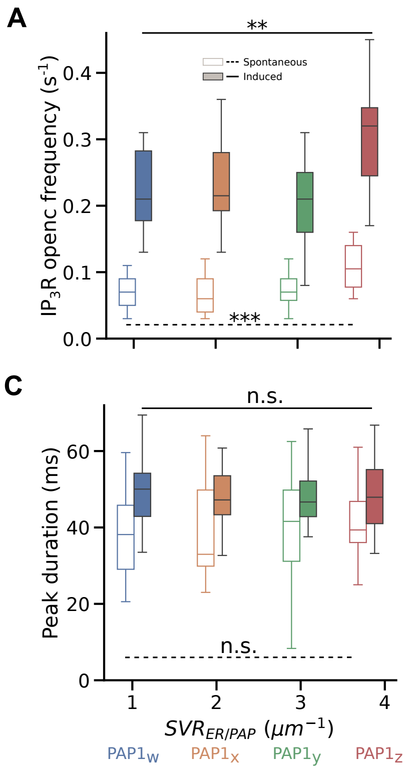


Fig S13: $Ca^{2+}$ **peak frequency increases as** $SVR_{ER/PAP}$ **increases independently from the number of** $IP_3R$ **channels on the ER.** In Fig. 4, the total number of $IP_3R$ channels, $N_{IP3R}$, varied depending on the mesh. Here, simulations were performed with the same number of $IP_3R$ channels on the ER of all meshes: 460 (same as $PAP1_z$ in Fig. 4). (A-D) Quantification of $IP_3R$ opening frequency (A), $Ca^{2+}$ peak frequency (B), duration (C) and amplitude (D) in $PAP1_w$,$PAP1_x$, $PAP1_y$, and $PAP1_z$, which vary in $SVR_{ER/PAP}$ and median $d_{ER-PM}$. $IP_3R$ opening frequency (ANOVA, $p = 4.63 \times 10^{-4}$ and $p = 7.67 \times 10^{-3}$ for spontaneous and induced signals, respectively) and $Ca^{2+}$ peak frequency (ANOVA, $p = 0.012$ and $p = 0.031$ for spontaneous and induced signals, respectively) increase as $SVR_{ER/PAP}$ increases. However, $SVR_{ER/PAP}$ had no significant effect on either spontaneous nor induced peak duration (ANOVA, $p = 0.88$ and $p = 0.84$, respectively) and amplitude (ANOVA, $p = 0.93$ and $p = 0.92$, respectively).

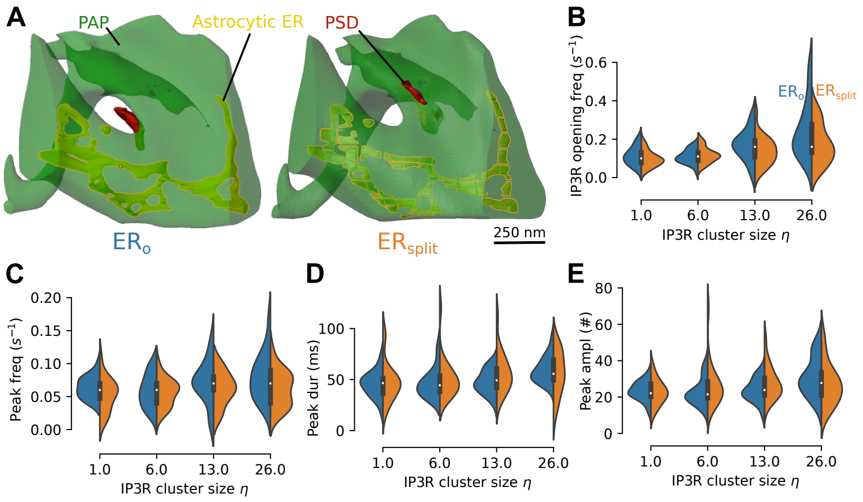


Fig S14: **Splitting the ER does not alter $Ca^{2+}$ activity in the PAP mesh.** (A) Screenshots presenting the PAP mesh from synapse d1s15a32b1 before (left, 'ER$_0$') and after (right, 'ER$_{split}$') performing the ER splitting step of the automated mesh generation pipeline presented in Fig. 5A. (B-E) Quantification of IP$_3$R opening frequency (B), free $Ca^{2+}$ peak frequency (C), duration (D) and amplitude (E) for IP$_3$R cluster size $\eta$=1, 6, 13 and 26, in 'ER$_0$' (left, blue) and 'ER$_{split}$' (right, orange) meshes. No significant difference was observed between $Ca^{2+}$ peak characteristics in 'ER$_0$' and 'ER$_{split}$', for all values of IP$_3$R cluster size tested.

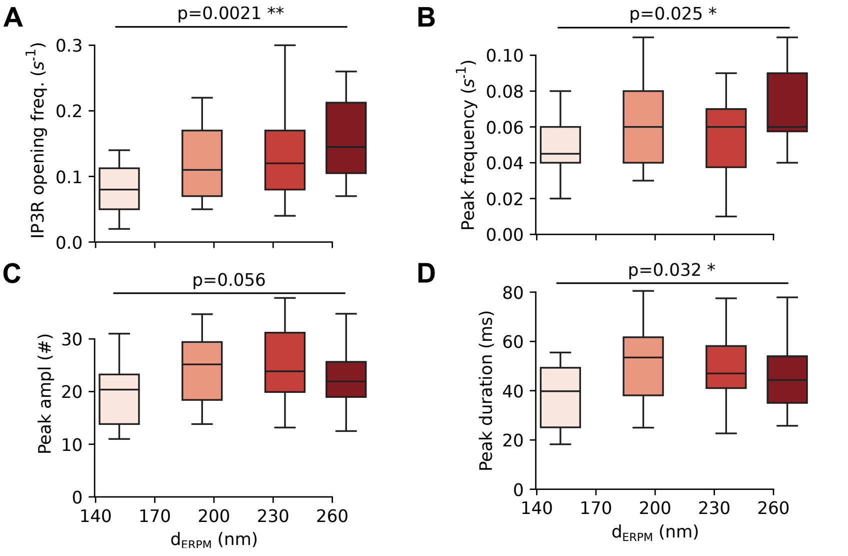


Fig S15: $Ca^{2+}$ **microdomain activity varies with** $d_{ERPM}$. To test whether the effect of $d_{ERPM}$ on $Ca^{2+}$ activity observed in mesh d1s15a32b1 was reproducible in other PAP meshes, the algorithm presented in Fig. 5A was used to create 3D PAP meshes derived from mesh d9s4a34b1 (presented in Table 1 and Figure 3) with various $d_{ERPM}$ and constant shape, volume, and surface area of PAP and ER. $IP_3R$ opening frequency (A), $Ca^{2+}$ peak frequency (B), amplitude (C), and duration (D) increased with $d_{ERPM}$.

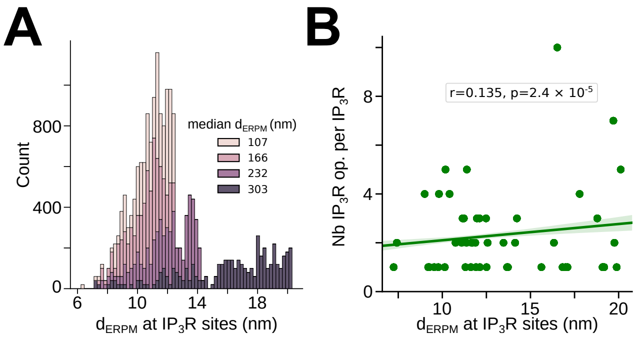


Fig S16: $d_{ERPM}$ **at $IP_3R$ sites dictates $IP_3R$ opening frequency.** . (A) $d_{ERPM}$ at $IP_3R$ sites increases with median $d_{ERPM}$. (B) The number of opening events of individual $IP_3R$s increases with $d_{ERPM}$ at the $IP_3R$ site.

#### Supplementary Movie 1

**Rendering of the reconstructed hippocampal astrocytic volume and the neighbouring synapses.** Rendering of the 220 $\mu m^3$ (7.07 $\mu$m x 6.75 $\mu$m x 4.75 $\mu$m) hippocampal astrocytic volume from the CA1 stratum radiatum region reconstructed from a perfectly isotropic electron microscopy stack (6 nm voxel resolution), together with the twenty seven fully reconstructed synapses in contact with the astrocyte (see Fig. 1 and Methods for details), zooming in on the d1s9a60b1 synapse. Tripartite synapse meshes contain the following elements: the perisynaptic astrocytic process (PAP, green), the astrocytic endoplasmic reticulum (ER, yellow), the bouton (grey) and the spine (blue).

#### Supplementary Movie 2

**Visualization of a simulation in a realistic PAP mesh extracted from electron microscopy** Movie illustrating a simulation in the PAP mesh from synapse d9s4a34b1. $IP_3$ (red) and Ca-GCaMP molecules (yellow) diffuse in the cytosol. $IP_3R$ channels (blue) are located at the ER membrane and $Ca^{2+}$ channels $Ch_{PM}$ (purple) are located on the plasma membrane. Molecule position is updated every 0.01 ms. Note that the size of molecules in the movie is increased for visualization purposes and that the darker and lighter greens result from 3D shading and rendering of the meshes.

#### Supplementary Movie 3

**Automatic redistribution of split ER in a realistic 3D PAP mesh reconstructed from EM.** Movie of a simulation of n frames generated in Blender, which alters the location of the ER objects within the PAP (here, d1s15a32b1). Each frame is thus characterized by a unique distribution of the ER objects within the PAP, while ER and PAP shape, surface area, volume and SVR are constant across frames.

#### Supplementary Movie 4

**Visualization of a simulation in a realistic PAP mesh extracted from electron microscopy at stimulation time.** Movie illustrating a simulation in the PAP mesh from synapse d2s6a9b1, at neuronal stimulation time: t=1s. 50 $IP_3$ molecules (red) are injected in tetrahedra below the plasma membrane of the PAP, emulating $IP_3$ synthesis resulting from the activation of metabotropic glutamatergic receptors at the membrane of the PAP. $IP_3$ and Ca-GCaMP molecules (yellow) diffuse in the cytosol. $IP_3R$ channels (blue) are located at the ER membrane and $Ca^{2+}$ channels $Ch_{PM}$ (purple) are located on the plasma membrane. Molecule position is updated every 0.1 ms. Note that the size of molecules in the movie is increased for visualization purposes.

### References

[1] C. Calì, M. Wawrzyniak, C. Becker, B. Maco, M. Cantoni, A. Jorstad, B. Nigro, F. Grillo, V. D. Paola, P. Fua, and G. W. Knott, "The

effects of aging on neuropil structure in mouse somatosensory cortex—A 3D electron microscopy analysis of layer 1," *PLOS ONE*, vol. 13, p. e0198131, July 2018. Publisher: Public Library of Science.

[2] A. Denizot, M. Arizono, U. V. Nägerl, H. Soula, and H. Berry, "Simulation of calcium signaling in fine astrocytic processes: Effect of spatial properties on spontaneous activity," *PLOS Computational Biology*, vol. 15, p. e1006795, Aug. 2019.

[3] A. Denizot, M. Arizono, U. V. Nägerl, H. Berry, and E. De Schutter, "Control of Ca2+ signals by astrocyte nanoscale morphology at tripartite synapses," *Glia*, vol. 70, no. 12, pp. 2378–2391, 2022. _eprint: https://onlinelibrary.wiley.com/doi/pdf/10.1002/glia.24258.
